# Functional similarity of non-coding regions is revealed in phylogenetic average motif score representations

**DOI:** 10.1101/2023.04.09.536185

**Authors:** Aqsa Alam, Andrew G Duncan, Jennifer A Mitchell, Alan M Moses

## Abstract

Here we frame the cis-regulatory code (that connects the regulatory functions of non-coding regions, such as promoters and UTRs, to their DNA sequences) as a representation building problem. Representation learning has emerged as a new approach to understand function of DNA and proteins, by projecting sequences into high-dimensional feature spaces, where the features are learned from data by a neural network. Inspired by these approaches, we seek to define a feature space where non-coding regions with similar regulatory functions are nearby each other. As a first attempt, we engineered features based on matches to biochemically characterized regulatory motifs in the DNA sequences of non-coding regions. Remarkably, we found that functionally similar promoters and 3’ UTRs could be grouped together in a feature space defined by simple averages of the best match scores in (unaligned) orthologous non-coding regions, which we refer to as phylogenetic average motif scores. Perhaps most important, because this feature space is based on known motifs and not fit to any data, it is fully interpretable and not limited to any particular cell type or experimental context. We find that we can read off known regulatory relationships and evolutionary rewiring from visualizations of phylogenetic average motif score representations, and that predicted regulatory interactions based on neighbors in the feature space are borne out in transcription factor deletion experiments. Phylogenetic averages of match scores to known motifs is a baseline for representation learning applied to non-coding sequences, and may continue to improve as databases of motifs become more complete.

## Introduction

Understanding how the DNA sequences of non-coding regions determine gene expression changes and cell-type specific expression patterns remains a central challenge in genome analysis; it is referred to as the cis-regulatory code [1–3]. Key to understanding the cis-regulatory code are sequence specific transcription factors that control the rate of transcription initiation and elongation [1]. In the past two decades, increasingly sophisticated experimental techniques have been developed to identify the targets of transcription factors in cells [4–6], and these data have been integrated to produce functional genomics maps at the genome scale[7,8]. Another regulatory code is believed to control mRNA stability, localization and splicing [9,10], and there are analogous techniques for high-throughput interrogation of RNA-binding proteins that control these processes[11,12]. However, even when integrated, multi-omics data appear to be less useful than anticipated in predicting transcriptional regulatory networks [1,13,14] and inferring strongly predictive regulatory codes from these data remains a challenge [13].

Recently, deep learning (and other advanced computational) methods trained on high-throughput cellular assays have been shown to predict gene expression directly from non-coding DNA sequences [15–18], suggesting that an understanding of the cis-regulatory code might be within reach[19,20]. Extending these approaches to generalize beyond the cell-types and data on which they were trained is now a major goal [21,22]. These methods learn directly from data to use large numbers of features and combinations thereof, but it is not yet clear whether it is the sheer number of comparably scaled informative features learned by these methods, the quality or novelty of the features, or their non-linear combination that is responsible for the predictive success. Studies from other areas of deep learning often reveal that it is the quality of the dataset specific features [23] and their appropriate scaling [24,25] that leads to the improved performance, rather than the non-linear combinations of features in higher layers of the encoder (e.g., [26]). Indeed, genomics deep learning approaches often appear to learn known DNA motifs as features[15,27,28] and in some systems, at least some aspects of regulatory function can be predicted using simple combinations of motifs, with relatively few constraints on their order orientation or distance [19,29–31].

In parallel to developments in computational methods, databases of genome sequences [32] and consensus motifs (Position Weight Matrices, PWMs) for RNA-binding proteins and transcription factors have continued to expand [33–37]. Databases of experimentally determined transcription factor motifs now reflect a substantial fraction of the “vocabulary” of the cis-regulatory code[37]. Although it is widely appreciated that transcription factor target genes cannot be reliably predicted from matches to their consensus motifs (the so-called futility theorem[38]), evolutionary conservation of motif-matches has been shown to increase the power of motif-matching[39–41]. Unfortunately, alignment-based motif-conservation approaches are unable to leverage the large numbers of genome sequences now available: they are limited by errors in sequence alignments of highly diverged non-coding sequences [42], loss of synteny of binding sites due to so-called “turnover” [43–47], as well as the technical difficulty of scaling phylogenetic analysis to large numbers of species (see, e.g., [48,49]). A further potential issue is that transcriptional regulatory networks are not always conserved between species[50,51].

Here we aim to reframe the problem of understanding how the function of non-coding regions is encoded in their DNA sequences as a representation building problem. We seek a feature space where non-coding DNA sequences with similar biological functions are nearby each other. Searching for similar non-coding regions in this space would allow us to predict function, analogous to the way BLAST uses sequence similarity to identify proteins of similar function. If we are successful in building a representation that captures *global* similarity of biological function, our predictions will immediately generalize across cell types and experimental conditions[21]. As a first step, we tested whether matches to experimentally determined PWMs could be used to build a functionally informative representation. To incorporate evolutionary information, we treat the motif match score as a quantitative trait and simply average over sets of orthologous non-coding sequences, similar to other studies that used simple summaries of motif matches in orthologs[52–54]. Although this approach is expected to have less power than rigorous phylogenetic approaches [40,41] and does not capture spacing, number and strength of motif matches as is possible with motif-based deep learning models[18,30,55], it has the clear advantages of no need for training data in any specific condition, and scaling to large numbers of homologs with no need for phylogenetic inference or tunable parameters of any kind.

Remarkably, despite the simplicity of our approach (no deep learning, no parameter fitting, no training data, etc.), we found that, from sequence alone, we were able to recall known functions and predict new functions of promoters in *S. cerevisiae* and, to some extent, human. Similarly, when we build a representation for 3’UTR sequences of *S. cerevisiae* using RNA-binding protein motifs, patterns in the representation space are strongly associated with sub-cellular localization of mRNAs. Our results suggest that given the relevant motifs, prediction of function for non-coding DNA might be easier than currently believed.

## Results

### Building a phylogenetic average motif score representation for non-coding regions

In the computer science literature, “feature extraction” (reviewed in [56]) refers to defining a collection of mathematical functions that map input such as images or human language to a feature space or “representation” that can be used for machine learning. Ideally, these features are designed so that semantically similar objects are grouped together. In bioinformatics, decomposing DNA sequences into k-mers (or gapped k-mers) is the most common way to extract putatively relevant features for non-coding regions (reviewed in [57]) although many other approaches have also been considered (e.g., [58]).

Here we set out to define a collection of knowledge-based features that could be extracted directly from DNA sequences to create a functionally informative representation vector for a non-coding region of interest. In our representation, each motif (Position Weight Matrix, PWM) denotes a feature, such that a representation built with k PWMs becomes a k-dimensional representation. Because PWM matches in individual DNA sequences occur frequently by chance [38], we averaged the (maximum) PWM scores over homologs (see Methods). For example, the Hsf1 transcription factor binding motif (TFBM) returns high match scores in the *S. cerevisiae* promoter sequences *HSP26*, which is regulated by Hsf1[59], and *RKM3*, which is not regulated by Hsf1 (Figure 1A). Clearly, high match scores alone in promoter sequences are not an indication of regulation. However, taking orthologous sequences into account, the maximum match scores for Hsf1 in all the orthologs of *RKM3* are significantly lower than the match scores for Hsf1 in the orthologs of *HSP26* (paired t-test: t = 4.4, p-value = 0.0007, n = 14) (Figure 1A). We summarize this by simply taking the average of the match score for all orthologs for each PWM, and assigning these scores to the non-coding regions of interest: in this example, for Hsf1, the *RKM3* promoter would get an average maximum motif score of 6.2 and the *HSP26* promoter would get 9.9 (Figure 1A, mean of orange points vs. mean of blue points, respectively). By averaging the match scores for motifs across evolution (which we refer to as phylogenetic averaging), we can prevent potential false matches in non-coding sequences from obscuring the real biological signal.

**Figure 1:**
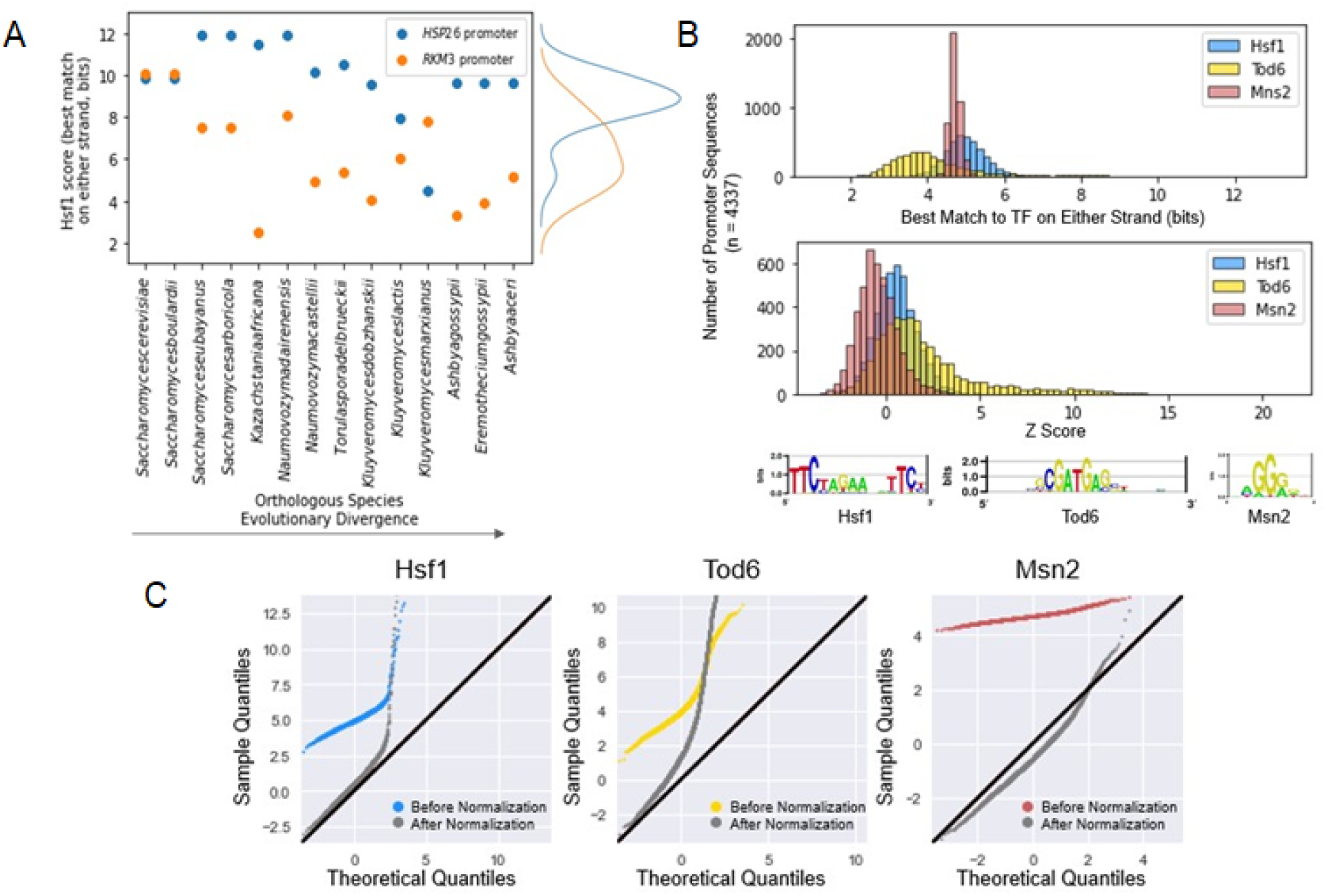
Overview of computational method. A) Yeast Hsf1 PWM scanned against *HSP26* and *RKM3* promoters in *S. cerevisiae* and its orthologs. Blue and orange dots correspond to the best match score to Hsf1 on either strand for *HSP26* and *RKM3* respectively. The x-axis represents orthologous species ordered from left to right according to approximate evolutionary divergence. Density plot on the right represents distribution of the scores for both *HSP26* and *RKM3*. B) Top panel shows raw distribution of average best match scores for 3TFs (Hsf1, Tod6, and Msn2 in blue, white, and red respectively) and bottom panel shows scores after normalization (see text and Methods for details). The y-axis indicates the number of promoter sequences in each bin for a total of n = 4337 promoter sequences. C) QQ-plots testing the standard normality of the distributions in b) before and after normalization. In each panel, the coloured lines (blue, yellow, and red for Hsf1, Tod6, and Msn2 respectively) indicate the raw scores, gray indicates the normalized scores, and black indicates y = x.

We next sought to ensure that the distributions of (phylogenetic averaged maximum) scores for each motif are comparable to each other, so that we could use them as features in a high-dimensional representation space. Non-coding sequences vary in GC content, repetitive elements, and other biases that are expected to affect motif scores. For example, random background matches to GC-rich PWMs will have higher scores in GC-rich promoters. Further, PWM match scores depend on the length of the PWM and its information content, and therefore scores from different PWMs cannot be directly compared. To address this, we normalized our features by randomly scrambling the positions of the PWM 100 times (which preserves the GC-content and information content of the PWM) and repeating the scanning for each scrambled matrix, then computing a z-score between the real match scores and the distribution of the scrambled match scores (see Methods). The output of this normalization resulted in a representation vector whose values are the z-scores comparing the real motif to scrambled motif scores. To confirm the normalization of the distribution of motif scores, we looked at matches of *S. cerevisiae* promoters to three different PWMs: Hsf1, a relatively long motif; Tod6, a medium length motif; and Msn2, a relatively short motif. The raw distributions of the best match score averaged over orthologs for 4337 *S. cerevisiae* promoter sequences (See Methods) for these PWMs are not standard normally distributed, and so it is difficult to compare their distributions. After applying our normalization method, we found that the distributions of Z-scores for each PWM were more comparable (Figure 1B). In fact, we found that much of the distribution was approximately standard normal except for the deviation in the positive tail which we believe to be the biological signal of highly conserved high scoring motifs (Figure 1C). Thus, we were able to make the phylogenetic average scores of different PWMs comparable.

### Phylogenetic average motif score representations are strongly associated with biological function

We first sought to perform unsupervised analysis on the knowledge-based representations. We built two promoter representations: one for *S. cerevisiae* promoters and one for human promoters: 4337 *S. cerevisiae* promoters (500 bp upstream of the start codon of the gene) with 244 PWMs from YeTFaSCo [34] and 15,906 human promoters (700 bp upstream of the annotated transcription start sites of Ensembl canonical promoters[60]) with the 137 non-redundant JASPAR 2022 [33] core PWMs. We clustered and visualized the representation vectors using standard approaches (see Methods).

Remarkably, despite no “fitting” of any parameters or training data, we found that patterns of phylogenetic average motif scores in yeast promoters (Figure 2A) and to some extent human promoters (Figure 2B) were associated with gene function. Using the SGD GO term finder[61] and g:Profiler (Raudvere et al., 2019) respectively (see Methods), we discovered at least 26 different clusters significantly enriched for various GO categories in both our yeast (Table 1) and human (Table 2) promoter sequence representations (False Discovery Rate [FDR] = 0.05, Benjamini-Hochberg corrected). At least for yeast promoters, the functional enrichments for these clusters, based only on sequence analysis, are comparable to those obtained by classic clustering of gene expression patterns[62], even though no gene expression measurements were used in our approach.

**Figure 2:**
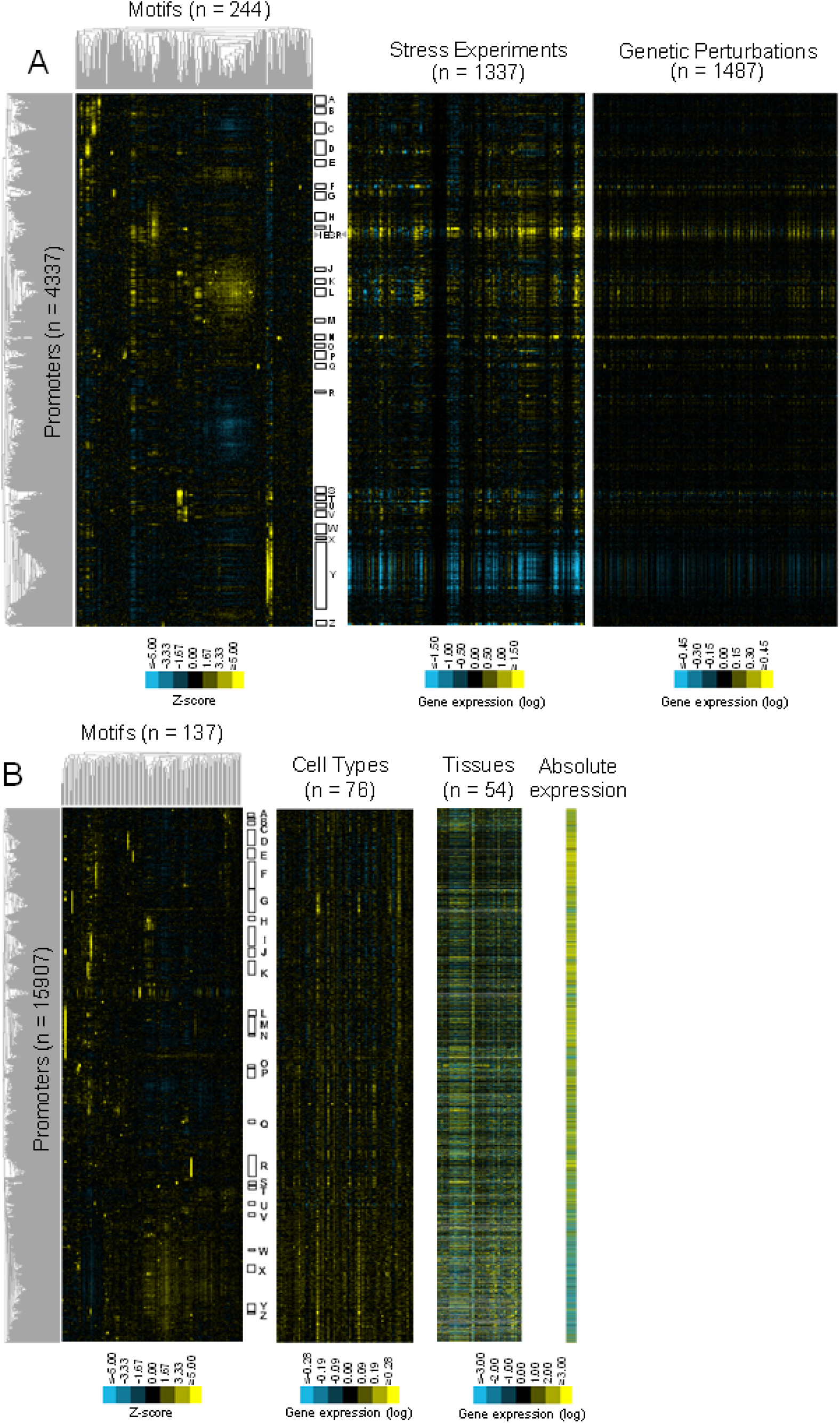
Clusters in the phylogenetic average motif score feature space are associated with biological function. A) The left heatmap shows the sequence-based representation of yeast promoter sequences, where each row is a promoter (total indicated by n) and each column is a known motif (total indicated by n). Bright yellow squares indicate motifs that on average have much higher scores in homologous promoters than scrambled versions of those motifs. Center and left panels show gene expression measurements in stress conditions and genetic perturbations (total experiments indicated by n), ordered by the rows of the clustering of the sequence-based representation (left) heatmap. iESR indicates a cluster of promoters with clear expression similarity, but no GO function association. B) human promoter representation compared to cell type-and tissue-specific gene expression patterns, as well as absolute RNA levels obtained from RNA-seq data. In the left panels, dendrograms represent the hierarchical clustering of the promoters (left) and the motifs (top). Letters represent enriched annotations (see Table 1 for yeast and Table 2 for human).

**Table 1:** Functional GO enrichments of clusters in the yeast promoter representation (SGD GO Term Finder)

| ID | PWMs | Annotations (Positive genes in cluster/Total genes in cluster) | Corrected P-value <= |
| --- | --- | --- | --- |
| A | Reb1, Stb2, Rtg3, Tye7, Mbp1, Rsc3, Swi4, Hcm1, Fkh1, Fkh2 | Microtubule binding (12/44), mitotic cell cycle (32/44), chromosome (18/44), microtubule cytoskeleton (19/44), chromosome segregation (25/44) (23/44) | 5.47e-16 |
| B | Nsi1, Reb1, Stb2, Pho4, Rtg3, Tye7, Abf1, Mbp3, Swi4, Rsc3, Fkh1, Fkh2 | Protein binding (20/49), mitotic cell cycle (18/49), cell division (15/49), cytoskeleton (14/49), cell cycle process (23/49) | 9.76e-06 |
| C | Stb2, Mbp1, Rsc3, Swi4, Fhl1, Fhk2 | DNA binding (47/105), DNA repair (50/105), cellular response to DNA damage stimulus (52/105), chromosome (59/105), replication fork (22/105) | 2.37e-21 |
| D | Ino2, Pho4, Rtg3, Tye7, Cbf1, Met32, Met31 | Catalytic activity (23/29), sulfur amino acid metabolic process (14/29), sulfur compound metabolic process (18/29), sulfur amino acid biosynthetic process (12/29), methionine biosynthetic process (10/29) | 2.61e-05 |
| E | Nsi1, Reb1, Stb2 | Transport (27/49), RNA localization (10/49), establishment of localization (27/49), macromolecule localization (22/49), protein-containing complex (35/49) | 3.13e-05 |
| F | Hap4 | Proton transmembrane transporter activity (24/29), proton transmembrane transport (23/29), inner mitochondrial membrane protein complex (26/29), monatomic cation transmembrane transporter (24/29), mitochondrial protein-containing complex (26/29) | 3.24e-34 |
| G | Rpn4, Abf1, Reb1 | Proteasome complex (34/67), endopeptidase complex (34/67), ATP hydrolysis activity (15/67), modification-dependent protein catabolic processes (42/67), proteolysis involved in protein catabolic processes (43/67) | 8.02e-10 |
| H | Nsi1, Reb1, Stb2 | Actin filament binding (6/69), Arp2/3 complex-mediated actin nucleation (7/69), cortical cytoskeleton (11/69), cell cortex (14/69), actin nucleation (7/69) | 7.57e-05 |
| I | Yap5, Yap3, Yap7, Yap1, Cad1 | Oxidoreductase activity (15/18), antioxidant activity (8/18), response to oxidative stress (11/18), cellular oxidant detoxification (8/18), cellular detoxification (8/18) | 8.41e-13 |
| J | Rtg3, Gcn4, Vhr1, Vhr2, Bas1, Rtg1, Yap7, Lys14 | Carboxylic acid transmembrane transporter activity (5/12), lysine biosynthetic process via aminoadipic acid (7/12), lysine biosynthetic process (7/12), lysine metabolic process (7/12), aspartate family amino acid biosynthetic process (7/12) | 2.30e-06 |
| K | Various TFs | Transporter activity (12/21), xenobiotic transport (5/21), plasma membrane (10/21), active transmembrane transporter activity (9/21), xenobiotic export from cell (4/21) | 5.37e-07 |
| L | Various TFs | Catalytic activity (40/51), oxidoreductase activity (16/51), monocarboxylic acid metabolic process (23/51), fatty acid metabolic process (17/51), peroxisome (17/51) | 1.07e-07 |
| M | Uga3, Ume6, Stb5 | Meiosis I (12/21), meiotic cell cycle (16/21), meiotic nuclear division (13/21), condensed nuclear chromosome ((8/21), nuclear chromosome (10/21) | 4.93e-10 |
| N | Hsf1, Abf2, Sfl1 | Chaperone binding (9/22), unfolded protein binding (10/22), heat shock protein binding (7/22), protein folding (17/22), protein refolding (9/22) | 2.82e-11 |
| O | Upc2 | Sterol biosynthetic process (6/12), steroid biosynthetic process (6/12), sterol metabolic process (6/12), lipid biosynthetic process (8/12), ergosterol biosynthetic process (5/12) | 5.86e-08 |
| P | Mcm1, Fkh1, Fkh2, Mbp1, Rsc3, Swi4 | Single-stranded DNA helicase activity (5/39), Cell cycle (22/39), MCM complex (5/39), Nuclear pre-replicative complex (6/39), cell cycle process (19/39) | 7.60e-08 |
| Q | Ndt80, Sum1, Cup9 | Ascospore formation (22/39), cell development (22/39), spore (24/39), sexual sporulation (22/39), intracellular immature spore (12/39) | 1.50e-13 |
| R | Matalpha2-dimer, Ste12 | Conjugation with cellular fusion (11/14), sexual reproduction (11/14), mating projection tip (8/14), mating projection (8/14), reproduction (11/14) | 7.60e-09 |
| S | Rtg3, Gcn4, Vhr1, Vhr2, Bas1, Rtg1, Yap7 | Amino acid biosynthetic process (36/49), arginine biosynthetic process (8/49), glutamine family amino acid metabolic process (12/49), isoleucine biosynthetic process (7/49), threonine metabolic process (4/49) | 5.58e-06 |
| T | Rtg3, Gcn4, Vhr1, Vhr2, Bas1, Rtg1 | Aminoacyl-tRNA ligase activity (12/18), ligase activity forming carbon-oxygen bonds (12/18), tRNA aminoacylation for protein translation (12/18), amino acid activation (12/18), aminoacyl-tRNA synthetase multienzyme complex (3/18) | 4.39e-07 |
| U | Rtg3, Gcn4, Vhr1, Vhr2, Bas1, Rtg1, various Yap TFs | Carboxylic acid binding (4/17), organic acid binding (4/17), one-carbon metabolic process (5/17), small molecule metabolic process (13/17), glycine cleavage complex (3/17) | 1.46e-06 |
| V | Cad1, Yap7, Yap1, Rtg3, Gcn4, Vhr1, Vhr2, Bas1 | Iron-sulfur cluster binding (7/39), metal cluster binding (7/39), sulfur compound metabolic process (15/39), iron-sulfur cluster assembly (8/39), metallo-sulfur cluster assembly (8/39) | 2.53e-07 |
| W | Abf1, Stb3, Sum1, Sfp1 | Structural constituent of ribosome (26/95), translation (51/95), organellar ribosome (24/95), peptide biosynthetic process (51/95), mitochondrial ribosome (24/95) | 2.06e-15 |
| X | Tod6, Dot6, Stb3, Sfp1, Tho2 | RNA polymerase I core promoter sequence-specific DNA binding (2/21), RNA polymerase I core factor complex (2/21), RNA polymerase I transcription regulator complex (2/21) | 6.57e-03 |
| Y | Tod6, Dot6, Spt10, Stb3, Sum1, Sfp1 | Catalytic activity acting on RNA (100/491), RNA binding (175/491), ncRNA metabolic process (245/491), ribosome biogenesis (222/491), nucleolus (193/491) | 2.90e-30 |
| Z | Rap1 | Structural constituent of ribosome (19/30), cytoplasmic translation (16/30), translation (20/30), cytosolic ribosome (19/30), ribosomal subunit (20/30) | 2.39e-10 |

**Table 2:**
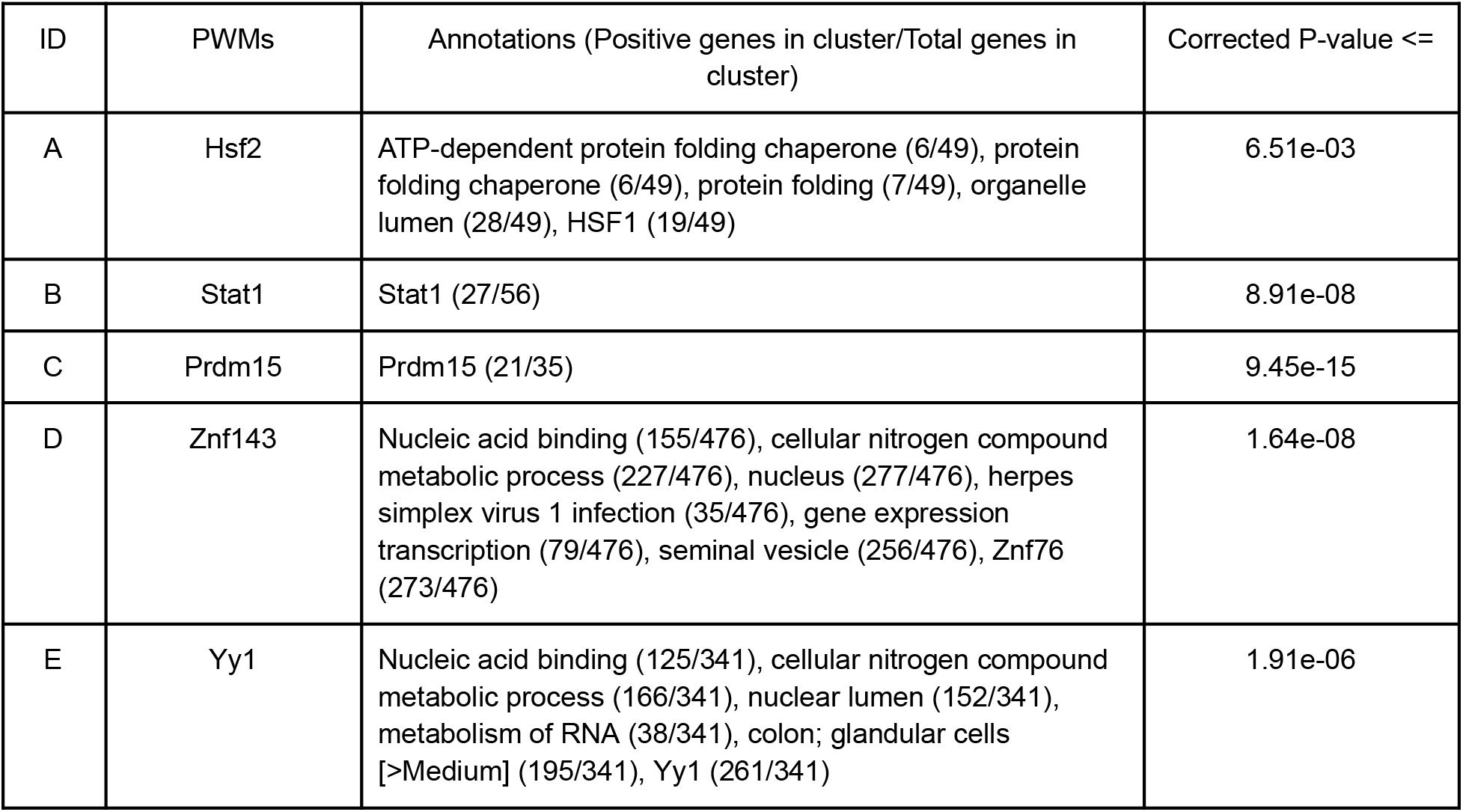
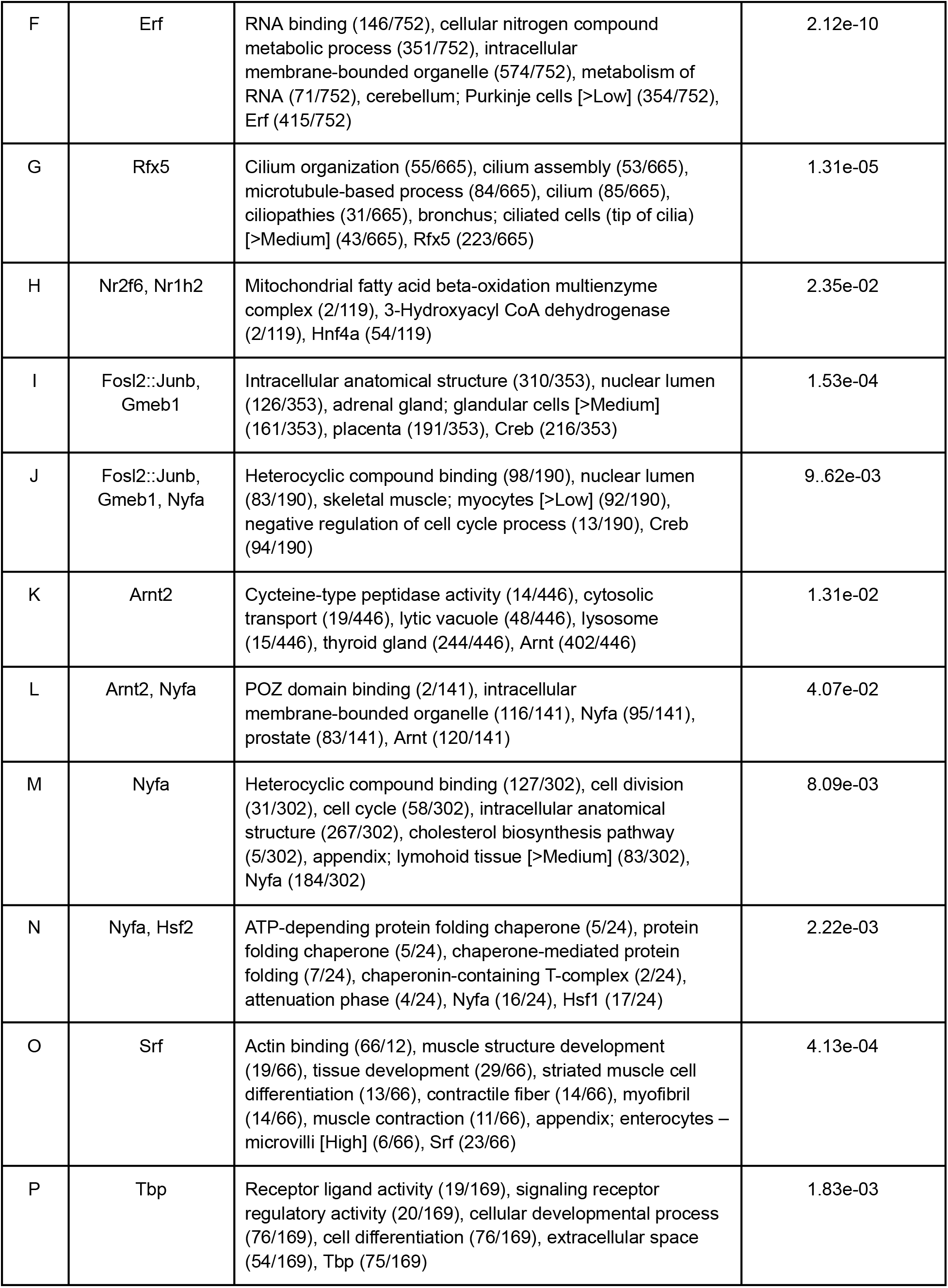
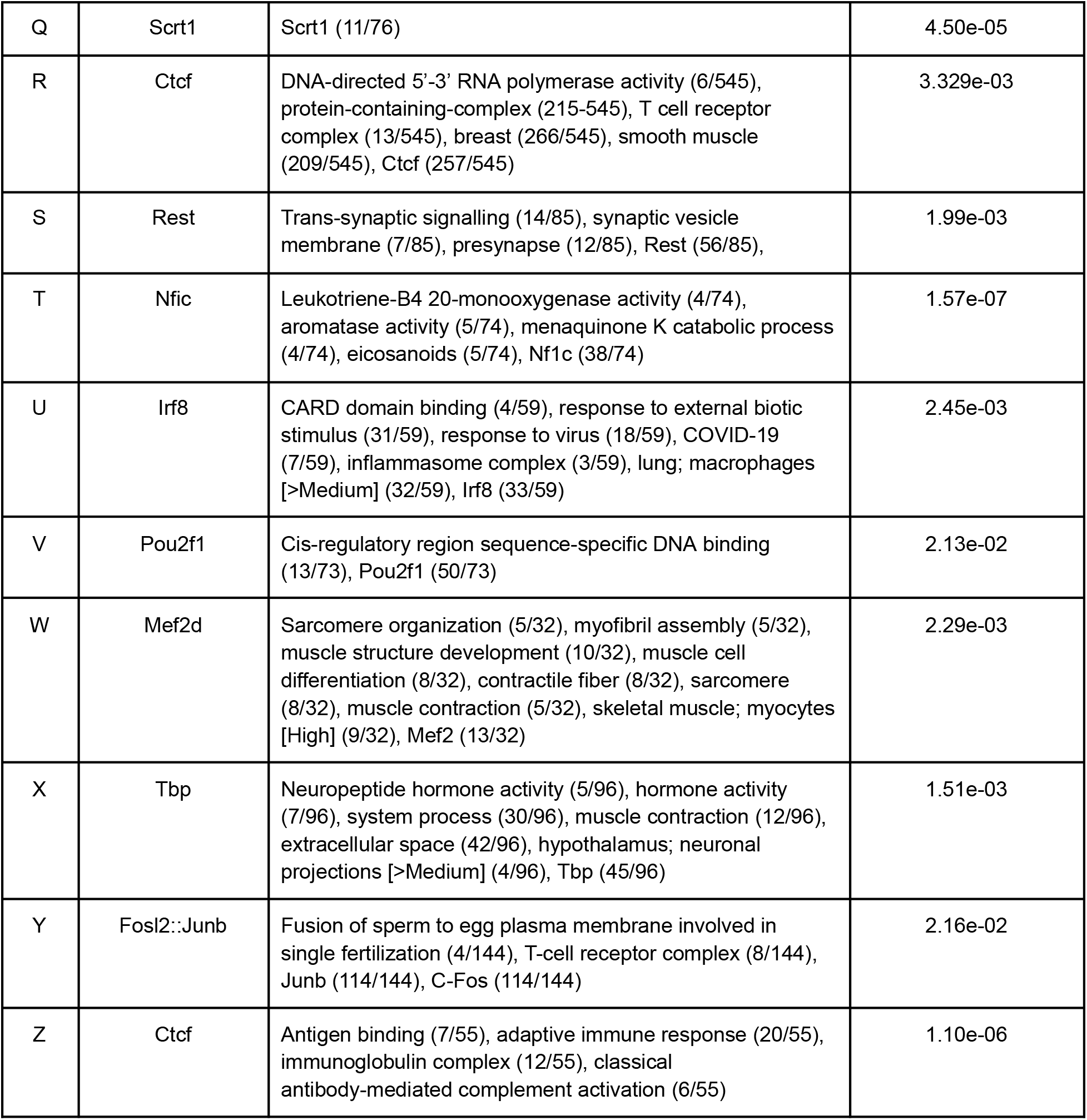
Functional GO enrichments of clusters in the human promoter representation (g:Profiler)

We next compared our yeast promoter representations to large-scale expression datasets (a compilation of >1000 stress experiments [63], and a collection of >1000 deletion experiments [64]) to see if the clusters of promoters we discovered in the sequence data drove similar gene expression patterns. We found broad agreement: clusters in phylogenetic average motif score space corresponded to genes with similar patterns of gene expression. We noted at least one cluster of yeast promoters (indicated below cluster I in Figure 2A) that share sequence signals, appear to drive similar expression patterns in many conditions (center panel in figure 2A), but were not enriched for any particular GO function. These promoters show signals for Msn2/4 and other STRE-like PWMs (Gis1, Com2) indicating that they are the induced stress-response promoters[65].

For the human promoter representation, the comparison with gene expression data (Human proteome atlas cell-type gene expression from sc-RNA-seq[66] and GTEx median expression, obtained from the GTEx Portal on 03/25/23) revealed a weaker association between gene expression patterns and cell-type and tissue-specific gene expression patterns (Figure 2B). Nevertheless, several of our sequence-based clusters showed clear cell-type specific gene expression patterns, For example, cluster S, which showed a strong signal for the neuronal repressor REST[67], showed clear expression in brain tissues. This level of tissue specificity was somewhat unexpected considering that enhancers were not included in the analysis, but is consistent with the finding that other sequence-based models are able to predict tissue-specific expression mostly using promoter sequences[22]. We also noted a clear association between the absolute expression level (median expression level across GTEx tissues) and the motif-based promoter representation (Figure 2B). Three of the clusters D, E, and R, which showed strong signals for Znf143, yy1 and CTCF, which are known regulators of enhancer promoter interactions[68,69], were associated with higher absolute expression levels. These results support the idea that human promoters contain a mixture of information about both tissue specific expression and absolute mRNA abundance.

To quantify the genome-wide association of the motif-based representation with gene expression, we assessed the power of the “guilt-by-association” approach in the representation space. We used nearest neighbour (NN) regression (also known as k-nearest neighbour regression with k=1, or 1-NN regression) to predict gene expression and summarized the predictive power of the regression by averaging the Pearson correlation over all the expression experiments in the datasets (numbers of experiments indicated in Figure 2). To avoid choosing a cutoff for similarity in this analysis, we required a prediction for each promoter under each condition, regardless of the similarity of the neighbour. Since there does not necessarily exist another promoter that drives an identical expression pattern for any gene, even an ideal representation is not expected to achieve a correlation of 1.0 in this analysis. Finding highly similar expression-pattern-driving promoters also becomes increasingly unlikely as the number of experimental conditions increase. Therefore, to obtain an upper bound, we computed the correlation of each gene with its nearest neighbour in the expression space. We also permuted the labels of our representation to estimate any correlation with gene expression measurements expected by chance. Consistent with the visual inspection (Figure 2) we found association with gene expression patterns (average R values around 0.2) far beyond what could be expected by chance (R values around 0), but still well below the upper bounds for these datasets (Tables 3 and 4). We note that these can be considered 0-shot sequence-based predictions of gene expression over diverse experimental contexts, a task that has proven difficult even for sophisticated deep learning models: in order to compare sequence-based models outside of their training data, other studies have fit regressions to part of the evaluation data[22].

**Table 3:** Functional GO enrichments of clusters in the yeast 3’UTR representation (SGD GO Term Finder)

| ID | Annotations (Positive genes in cluster/Total genes in cluster) | Corrected P-value <= |
| --- | --- | --- |
| A | Mitochondrial gene expression (132/430), mitochondrial translation (117/430), mitochondrion organization (132/430), mitochondrial part (132/430), mitochondrion (342/430), structural constituent of ribosome (79/430), mitochondrial ribosome (83/430) | 2.57e-37 |
| B | RNA biosynthetic process (104/273), transcription by RNA polymerase II (93/273), DNA binding (72/273), chromosome (78/273), nucleus (173/273) | 8.48e-17 |
| C | RNA binding (98/251), ribosome biogenesis (133/251), rRNA processing (100/251), nucleolus (116/251), preribosome (81/251) | 2.67e-19 |

**Table 4:** NN regression R values (Pearson correlation) for yeast promoter representation and gene expression data

| Experiment (number of expression conditions) | Randomized (average R) | Phylogenetic average motifs scores (average R) | Upper bound (average R) |
| --- | --- | --- | --- |
| Stress (n=1337) | 0.00267 | 0.23393 | 0.57255 |
| Deletions (n=1487) | -0.00051 | 0.17235 | 0.56796 |

**Table 5:**
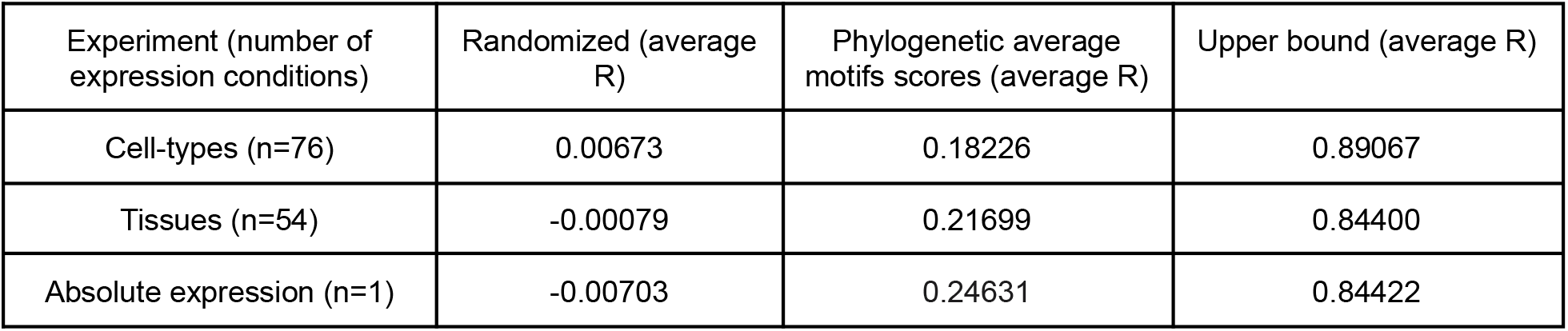
NN regression R values (Pearson correlation) for human promoter representation and gene expression data

| Experiment (number of expression conditions) | Randomized (average R) | Phylogenetic average motifs scores (average R) | Upper bound (average R) |
| --- | --- | --- | --- |
| Cell-types (n=76) | 0.00673 | 0.18226 | 0.89067 |
| Tissues (n=54) | -0.00079 | 0.21699 | 0.84400 |
| Absolute expression (n=1) | -0.00703 | 0.24631 | 0.84422 |

Because the strength of functional association with our sequence-based clusters was unexpected, we performed controls to rule out that simple primary sequence similarity was responsible. For example, we computed the pairwise percent identity (See Methods) between the promoters for three clusters of *S. cerevisiae* promoters and three clusters of human promoters that all showed strong associations with function (Figure 3). As expected from non-homologous promoter sequences, we found that all pairs show percent identity that is consistent with what can be observed in comparisons of randomly shuffled sequences (See Methods). This also confirms our expectation that functional similarity of non-coding sequences is generally not detectable using sequence alignments: the clustering of functionally similar promoters in the (sequence-based) phylogenetic average motif score representation space is remarkable.

**Figure 3:**
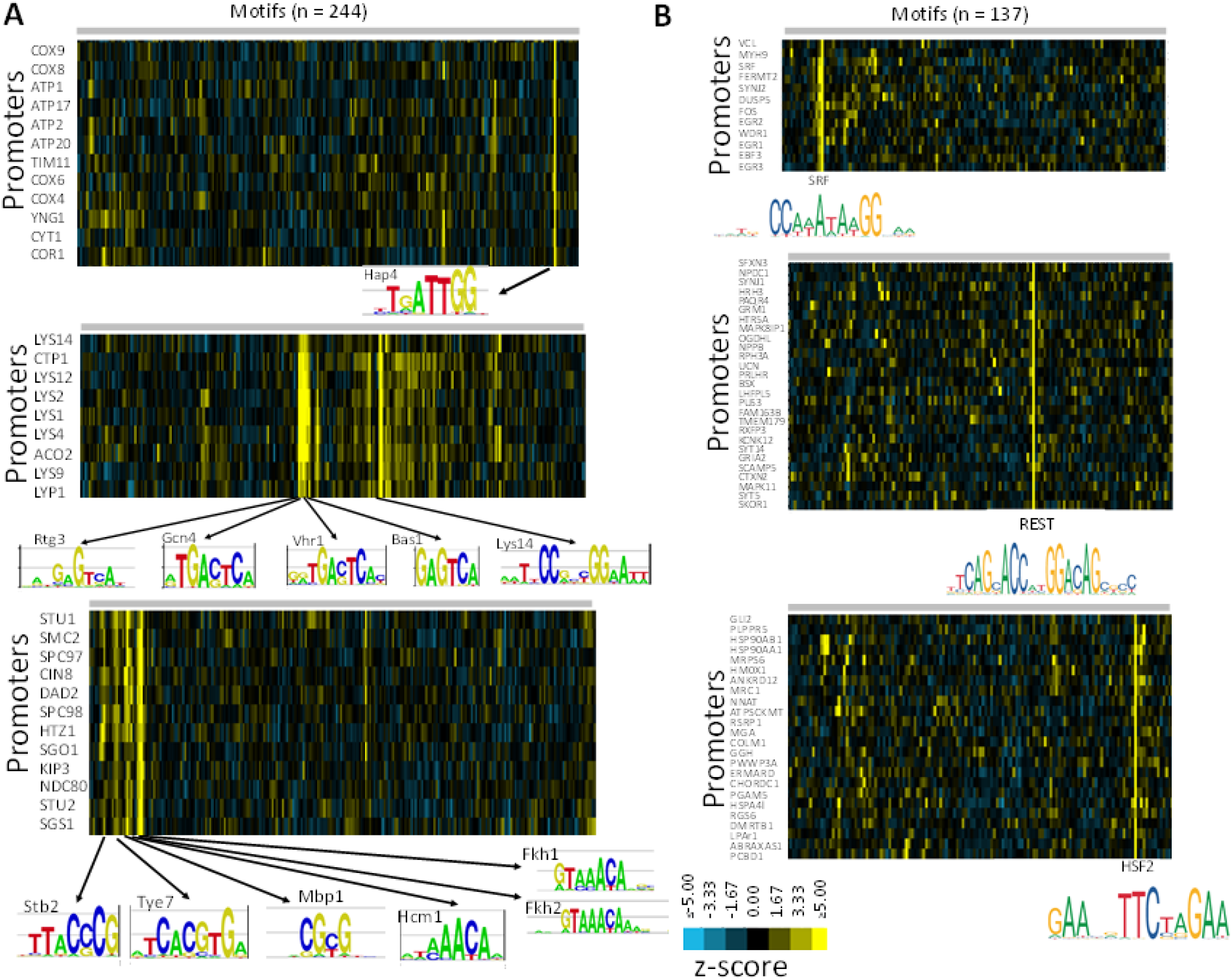
Diverse clusters reveal known regulators of biological processes in yeast and human promoters. Heatmaps of sequence-based promoter representation are displayed as in Figure 2. A) Clusters in the yeast promoter representation space associated with cytochrome c-oxidase and ATP synthase (top) lysine biosynthesis (middle) and chromosome segregation (bottom). Labels under each cluster represent the TFs with the highest z-scores in each cluster, with their logos (retrieved from YeTFaSCo). B) Clusters in the human promoter representation associated with striated muscle differentiation (top) trans-synaptic signaling (middle) and protein folding (bottom). Labels under each cluster represent the TFs with the highest z-scores in each cluster, with their logos (retrieved from JASPAR).

### Clusters in the phylogenetic average motif score representation for yeast and human promoters reveal known regulatory relationships and predict new ones

Because each feature in our representation corresponds to a transcription factor, we can predict regulatory relationships from the heatmap visualization. Interestingly, within the yeast promoter representation, we identify a few groups of promoters with strong signals for only one or two TFs. For example, Hap4 is known to regulate the expression of key mitochondrial complexes (Cytochrome c and ATP synthase) in the electron transport chain[70]. Remarkably, we find a cluster of promoters with a strong signal for only the Hap4 motif (Table1, cluster F), where 26/29 (90%) are annotated as belonging to protein complexes in the inner mitochondrial membrane. The other three promoters include *PMT2* and *YNG1*, both characterized genes that have no known association to respiration. Nevertheless, in this unusual case, it appears that phylogenetic average score for a single motif is a very strong predictor of promoter function. A slightly more complex example is the cluster of lysine biosynthesis promoters (Table 1, cluster J), where we find a clear signals for both the lysine specific factor, Lys14 [71], as well as the more general cellular amino acid biosynthetic regulator Gcn4 [72]. (We note that these promoters also contain strong signals for Rtg3, Vhr1, and Vhr2 which have similar PWMs to Gcn4 that likely cannot be distinguished in our analysis) Nevertheless, in these cases we can apparently read the cis-regulatory code directly from the visualization of the representation.

More typically, clusters of promoters show signals for multiple transcription factors. For example in the chromosome segregation promoters (Table 1, cluster A), a variety of TFs are highlighted: Hcm1, Fkh1/Fkh2, Tye7, and Stb2. While three of these (Hcm1, Fkh1, and Fkh2) have similar motifs, they are all known to regulate the cell cycle[73–75]. On the other hand, the roles of Tye7 and Stb2, (which have distinct motifs) in this process are unclear. However, Tye7 is cell-cycle regulated and Tye7 mutants are defective in sporulation[76]. Remarkably, Stb2 is not known to be involved in the yeast cell cycle, and this represents a new hypothesis of function for this transcription factor. However, we note that there is some similarity between the motif for Stb2 and the cell cycle regulators Swi4/6. Further experiments are needed to test how (or if) these factors work together to control transcription of chromosome segregation genes.

We identified other examples of strong signals for transcription factor motifs that represent new predictions for regulators of basic cellular functions. For example, the proteasome is known to be regulated by Rpn4[77] and a cluster that consists of mostly proteasomal promoters that all contain a strong signal for the Rpn4 PWM (Table 1 cluster G, supplementary Figure 1). However, in this cluster, we also find strong signals for Abf1 and Reb1, which to our knowledge have no reported role in proteasome regulation (Table 1 cluster G). Remarkably, Abf1 tends to have stronger signals in the promoters that tend to lack signals for Reb1 (Spearman’s R= −0.275, n=67 Z-scores for Abf1 and Reb1, *P*=0.025) and Reb1 and Abf1 have been previously reported to be interchangeable in at least one (unrelated) promoter[78]. We suggest a simple logic for these promoters: either Abf1 or Reb1 is needed in addition to Rpn4 in these promoters. Similarly, in addition to the known regulators of ribosome biogenesis (cluster Y; Stb3, Tod6/Dot6 and Sfp1) we observe a strong signal for Spt10, a factor that is not known to regulate the ribosome and has a low confidence motif not similar to the known factors. We speculate that either Spt10 is a new regulator of ribosomal biogenesis, or its motif is similar to another regulator (such as Ifh1) whose motif is not yet well-characterized.

Overall, we observed that most of the functions associated with phylogenetic average motif patterns include at least 3 different motifs, with additional, previously unreported motifs implicated for many functions. This suggests that for most yeast promoters, the full complexity of regulation is not fully understood. These observations are broadly consistent with the combinatorial nature of the cis-regulatory code [1].

Unlike the yeast promoter representation, the human promoter representation did not appear to have multiple motifs in each cluster; rather, each cluster was usually associated with only a single motif. Nevertheless, they reflected known regulatory relationships: we discovered clusters of genes involved in striated muscle differentiation, trans-synaptic signaling, and protein folding (Figure 3B). Interestingly, we noticed that the striated muscle differentiation cluster was associated with SRF and included the *SRF* promoter, which shows a strong signal for its own binding site. This is consistent with the known autoregulation of SRF[79]. Two other known SRF target genes, *FOS* and *ACTG2*[80] are in this cluster and contain strong signals for SRF. This demonstrates that we can capture some of the known biology using only phylogenetic averages of matches of TFs to promoter sequences. However, the small number of motifs associated with each cluster suggests that fewer or less diverse motifs are found in each human promoter (compared to yeast) or that a smaller subset of regulatory interactions are conserved during evolution.

### Testing predictions of regulation for yeast genes of unknown function

Having confirmed that known regulatory relationships were recovered in the phylogenetic average motif score representation, we next tested whether similarity in the representation space could predict the regulation of yeast genes of unknown function. We identified five genes of unknown function from diverse clusters and obtained gene expression patterns under relevant perturbations[61] for them and their neighboring genes in the sequence based clustering (Figure 4A-E). For example, we found YPR174C and the genes in its cluster all had increased mRNA expression after SWI4 was deleted, which is consistent with the expectations for DNA replication and repair genes regulated by Swi4[81] and with previous predictions that YPR174C is involved in DNA repair[82] (Figure 4A). As another example, we found that YKR005C contained a strong signal for UME6, which is known to regulate early meiosis genes and some middle stage meiosis genes, and YKR005C was reported to be a target of UME6 and is induced in later stages of meiosis[83] (Figure 4B). In all these cases, we were able to identify a relevant perturbation that affected expression of the genes in the sequence-based cluster, highlighting the ease of hypothesis testing using this fully interpretable knowledge-based representation.

**Figure 4:**
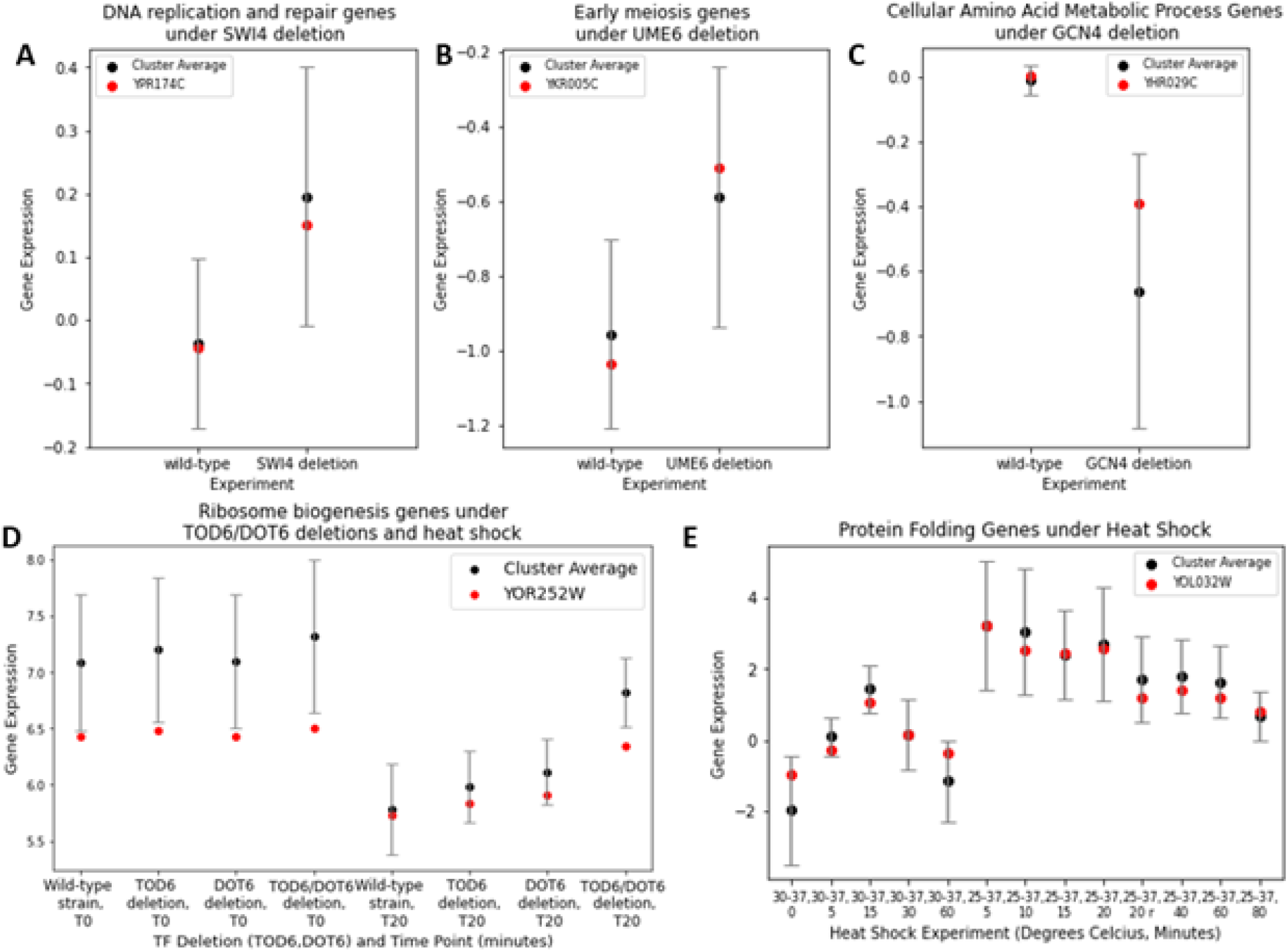
Predicting the regulation of unknown genes. The expression of genes of unknown function (red symbols) compared to their cluster (black symbols, clusters as defined in Table 1) under A) SWI4 deletion, B) UME6 deletion, C) GCN4 deletion, D) TOD6/DOT6 deletion and heat shock, E) heat shock. Gene expression data was obtained from the Saccharomyces Genome Database (SGD). Error bars on the black circles represent the standard deviation of the promoter sequence cluster mean, while red dots represent single value expression data points for a single gene of unknown function.

### A yeast 3’ UTR representation based on phylogenetic average RBP motif scores is associated with sub-cellular localization and binding of PUF proteins

Since our phylogenetic average motif score representation revealed functional similarity in promoter sequences, we next investigated if the same approach would lead to an informative representation for 3’ UTRs, which are implicated in regulation of mRNA localization and stability, but usually show little sequence similarity. We built a representation for 4294 *S. cerevisiae* 3’UTRs (defined as 200 bp downstream of the stop codon of the gene) using the 74 yeast RNA-binding protein motifs (RBPMs) from AtTract[35]. We found that clusters of 3’UTRs in this RBPM representation were associated with sub-cellular localization (Figure 5).

**Figure 5:**
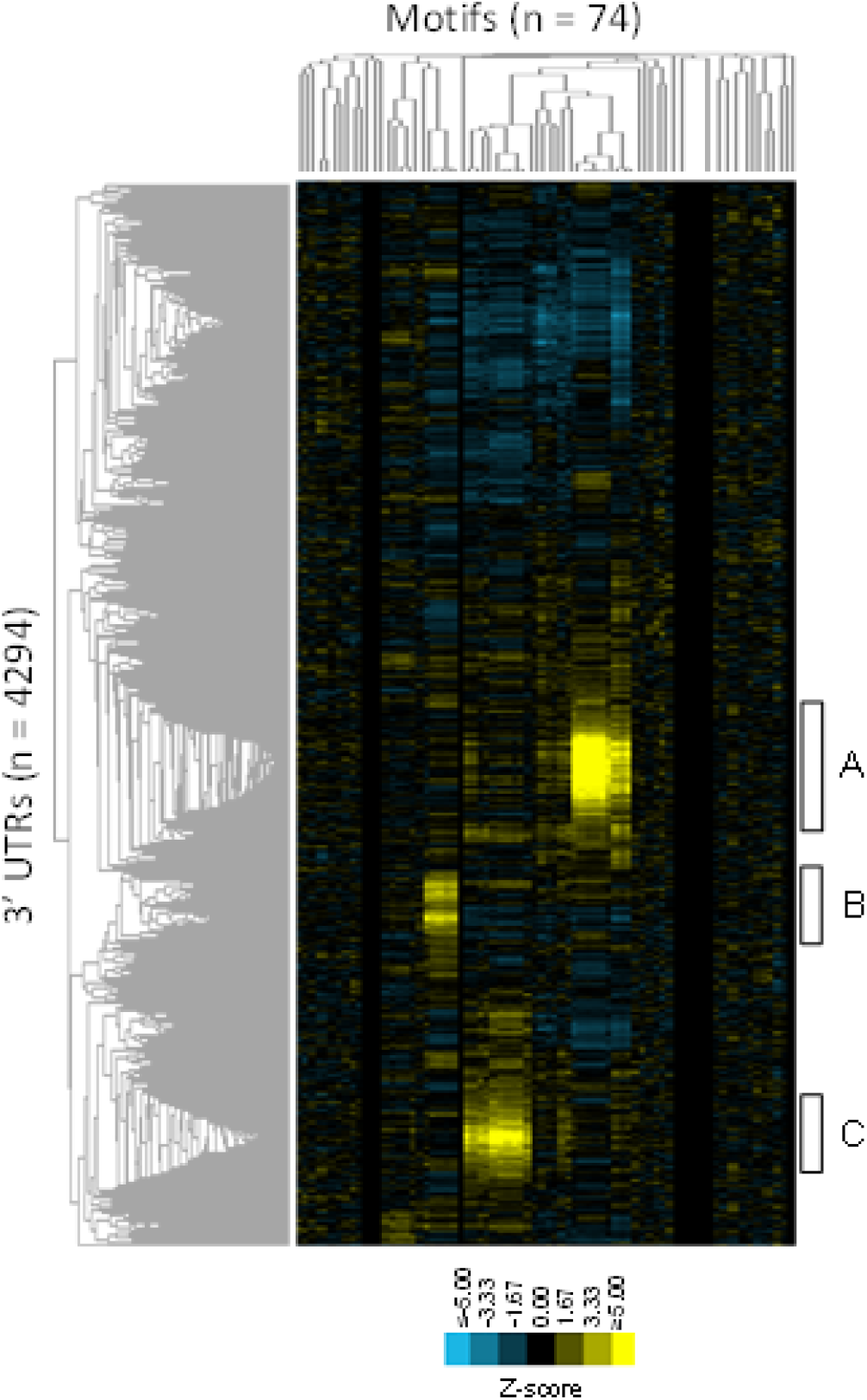
Functional similarity of yeast 3’UTRs. An phylogenetic average motif score representation for yeast 3’UTR sequences, displayed as for the promoter representations in figure 2. Letters correspond to enrichments described in Table 6. The dendrograms represent the hierarchical clustering of the UTRs(left) and the motifs (top).

**Table 6:** enrichment of GO localization and PUF binding in clusters of 3’ UTRs.

| ID | GO association positive in cluster/total genes in cluster (all p-value<1e-15) | Positive PUF3 in cluster/Total genes in cluster, (p-value) | Positive PUF4 in cluster/Total genes in cluster, (p-value) | Positive PUF5 in cluster/Total genes in cluster, (p-value) |
| --- | --- | --- | --- | --- |
| A | Mitochondrion 342/430 | **165/430 (2.92e-147)** | 10/430 (0.0261) | 10/430 (0.0810) |
| B | Nucleus 173/273, Chromosome 78/273 | 2/273 (2.16e-04) | 10/273 (0.131) | **36/273 (1.01e-14)** |
| C | Nucleolus 116/251, Preribosome 81/251 | 1/251 (8.51e-05) | **69/251 (7.59-45)** | 7/251 (0.150) |

We discovered 3 clusters significantly enriched for GO terms relating to localization to a cellular component: mitochondria, nucleus, and nucleolus (False Discovery Rate [FDR] = 0.05, Benjamani-Hochberg corrected) (Table 6). We found many fewer clusters compared to the promoter representations, perhaps due to the relative paucity and high redundancy of the RNA binding motif data (See Discussion). Nevertheless, we found that the association with mitochondrial localization was remarkably strong: 342/430 = 79.5% of promoters in this cluster drive expression of a nuclear encoded, mitochondrially localized protein(Table 6). This indicates that in yeast it is possible to predict mitochondrial protein localization from the 3’UTR of the mRNA, with positive predictive power of at least 80% for hundreds of transcripts. The appearance of these patterns in our representation is consistent with the idea that 3’UTRs are involved in localization of mRNA in the cell[84], and is similar to what was recently observed in a previous large-scale analysis of 3’ UTR evolution[54].

We noticed that each cluster seemed to be associated with a particular PUF motif, so we next checked for enrichment of PUF proteins in the clusters using in vivo RNA-protein binding data for PUF2, PUF3, PUF4, and PUF5 [85] in *S. cerevisiae*. We found that each of the three significant clusters was associated with a PUF binding protein (Table 6). Thus the phylogenetic average motif score representation appears to reflect the conserved binding of these well-characterized RNA binding proteins[53]. Interestingly, we noted that the association with GO annotated localizations are stronger than the association with experimental measurements of RNA protein binding, suggesting two (non-mutually-exclusive) possibilities: either there are other signals in the UTR sequences, or that the binding data are noisier than the phylogenetic average motif scores.

### Evolution is important for revealing biological signals despite transcriptional rewiring

Since the input to our encoder is a set of homologous non-coding sequences, we wondered how many orthologous species are required to reveal biological signals. Does including more species increase the biological signal, or does it plateau or decrease after a certain evolutionary divergence? To answer this question, we began with a fixed number of *S. cerevisiae* promoter sequences (n = 2925), each of which were associated with at least 17 orthologous sequences. We organized the orthologous species into 9 clades so we could increase the evolutionary divergence from one representation to another by including those species in groups (Figure 6A, black dots indicate presence of species). For each level of evolutionary divergence, we compute an phylogenetic average motif score representation as described above, and compared to gene expression using nearest neighbour regression as above. Since nearest neighbour regression has no trainable parameters and treats all dimensions of the representation equally, we can fairly compare our representation approach with standard sequence-based bioinformatics baselines: pairwise sequence similarity detected using BLAST (both promoter sequence and amino acid sequence of the encoded protein).

**Figure 6:**
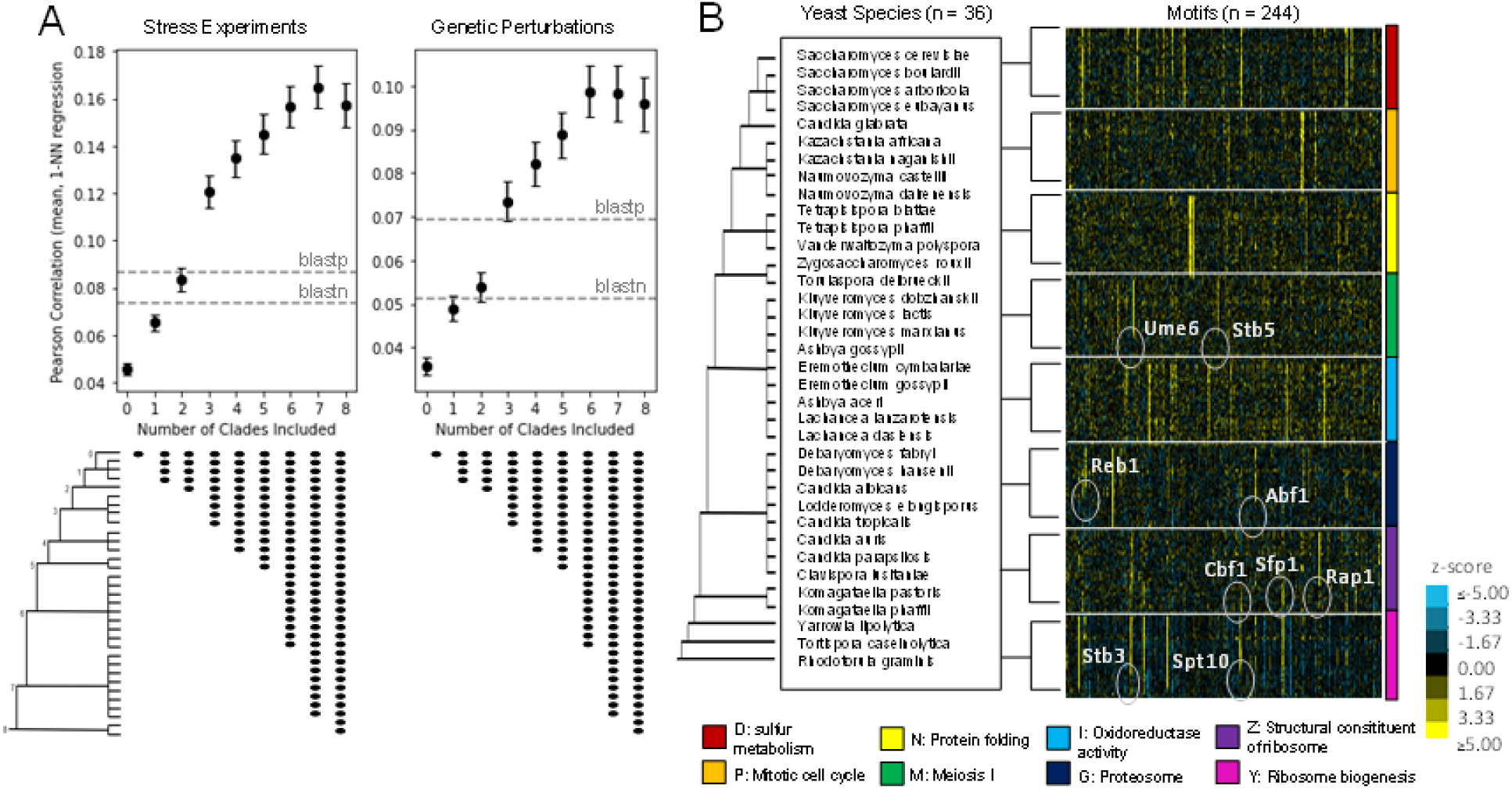
Evolution is important for detecting functional similarity of non-coding sequences. A) Shows the mean (filled symbol) and three times the standard error (error bars) of the Pearson correlation from nearest neighbour regression of gene expression based on similarity in the representation space (See Methods) as evolutionary divergence is increased. “Stress experiments” of 1337 stress-related conditions and “Genetic Perturbations” is a gene expression dataset with 1487 single gene deletions. The x-axis represents the number of clades included as depicted in the phylogenetic tree, and the y-axis represents the Pearson correlation. The branch lengths in the phylogenetic tree do not reflect evolutionary distances. Black dots under graphs represent species included at each evolutionary distance. B) Heat map depicting average motif scores in species in a particular functional cluster. Each square represents a z-score of the real motif score to a distribution of scores generated using 100 scrambled motifs averaged across a particular yeast species for genes in a particular cluster (See Methods). Clusters of *S. cerevisiae* promoters are described by the colours below the heat map and were extracted from Figure 2A. Within each cluster, species are organized roughly by evolutionary distance as described by the phylogenetic tree on the left. The dendrogram represents the hierarchical clustering of the motifs (top).

As expected, a representation based on the matches to the PWMs in *S. cerevisiae* alone showed less predictive power than BLAST. However, we found that as we increased the evolutionary divergence, the mean Pearson correlation increased, eventually surpassing the predictive power of BLAST, even if we used the amino acid sequence of the encoded proteins for BLAST (Figure 6A). We noted that there is a slight decrease or plateau in the Pearson correlation coefficient after 7 clades, suggesting that we had reached the limit of the useful divergence. We wondered whether the rewiring of the transcriptional network in distant species may be responsible for this plateau. To test this, we averaged PWM scores across the promoters for 8 of the clusters (as identified in Figure 2A) for each of the 36 species (Figure 6B) (see Methods). Remarkably, of these 8 examples, four showed conservation over regulation over nearly all the species here (Figure 6B) while for the other four the average signals for some motifs did not extend to all 36 species. This may explain the drop in predictive power observed in Figure 6A. Nevertheless, taken together, these results confirm that averaging over large numbers of orthologs was essential to the predictive power we observed in the yeast promoter representation (Figure 2).

Remarkably, we were able to visually identify examples of transcriptional regulatory rewiring from this analysis (indicated in gray on the heatmap in figure 6B). At least one of these is a known case, the transcriptional rewiring of ribosomal biosynthesis regulation [86–89]: Rap1 in *S. cerevisiae* to Cbf1 and Sfp1 in *C. albicans* and beyond (Figure 6B, purple). We also noticed that promoters of genes involved in Meiosis I appear to lose regulation by Ume6 (and Stb5, which has a similar motif, Figure 6B, green) in about half of the species considered here, which may be consistent with a previous report that predicted Ume6 as changing function in *C albicans* [90]. Similarly, while ribosomal biogenesis is regulated by Stb3 [91,92] in *S. cerevisiae*, this appears to be lost in the more distant species (Figure 6B, magenta). Two of the previously unreported associations we observed in *S. cerevisiae* (Abf1 and Reb1 in proteasomal promoters, dark blue, and Spt10 in ribosomal biogenesis promoters, purple) are also not found in distant species. Despite this transcriptional rewiring, we were able to group promoters of similar function together in the phylogenetic average motif score representation (Figure 2). We believe this is because the phylogenetic averaging retains signals that occur in a subset of species, albeit they appear quantitatively weaker (see Discussion).

## Discussion

Our effort to reframe the cis-regulatory code as a representation building problem was remarkably successful, especially considering the simplicity of the approach we took. To our knowledge, this is the first report of a DNA sequence-based measure of similarity for non-coding regions that is more informative than alignment-based sequence similarity using BLAST. Because the phylogenetic average motif score representations are based only on prior knowledge of motifs and homologous non-coding sequences, similarity of non-coding sequences in these spaces can predict function without relying on specific experimental data collected in a particular condition or cell type. Surprisingly, but consistent with recent results[13] and our finding that phylogenetic average motifs scores had stronger association with subcellular localization than with *in vivo* PUF protein binding data (Table 6), a representation of yeast promoters based on large-scale *in vivo* transcription factor binding data (192 transcription factors[93], See Methods) was a poor predictor of gene expression patterns (nearest neighbor regression, Pearson correlation, R = 0.003 +/- 0.006 and 0.004 +/- 0.02 on the yeast expression data sets described above).

The power of simple phylogenetic averaging of motif scores to predict gene expression is surprising because we know that many aspects of transcriptional regulation have changed during the hundreds of millions of years of divergence represented by the species considered here [51,52,90]. The naïve expectation is that if regulation is not conserved, this method should fail to identify functional similarity. However, in our example of yeast promoters, although the ribosomal transcriptional regulation is known to have switched from regulation by Tbf1 in *C. albicans* to regulation by Rap1 in *S. cerevisiae*[87], and both species were present in the representation, we were still able to identify a clear cluster of ribosomal proteins, and these were distinct from other promoters involved in translation (cluster W, Y and Z, Table 2). Additionally, the TFs identified as important (cluster W: Stb3, Sfp1, Y:Tod6, Dot6 and Z: Rap1, Table 2) were the expected TFs for each cluster. Thus, the approach can be quite robust to evolutionary changes in transcriptional regulation, perhaps due to our simple averaging of best match scores over all the orthologous species.

The high-dimensional phylogenetic average motif score feature space has the advantage of using all available TFs at once, which reflects the combinatorial nature of the cis-regulatory code[1]. Compared to deep learning approaches based on deep convolutional neural networks [27,94,95] or language models [28], the average motif score representation is trivially interpretable, and needs no downstream steps to identify important motif combinations (e.g., Abf1 and Reb1 working with Rpn4, supplementary figure S1). Because we can “read off” predictions from a visualization of the representation with no fine-tuning on additional datasets (as is needed for current deep learning approaches [22,28]), we can make highly specific biological predictions for cases where it would not be possible to create even a small training dataset for supervised analysis (e.g., Hap4 targets, Figure 3A). Indeed, for several genes of unknown function, we directly compared the predictions of regulation with TF knock out gene expression experiments (Figure 4).

One downside of our approach is that it relies on the quality of sequence and motif data. For example, the PWMs for RBPs are of lower quality and diversity than for TFs, and this may have limited the number of clusters we identified in the 3’ UTR analysis. Although we did not directly use any functional data to build the representation, there may still be some bias due to the way the motifs are derived: ChIP experiments are sometimes used to derive motifs[34]. Since these experiments are usually performed in standard lab conditions and cell lines, we may be missing motifs that would help predict other unknown functions. Motifs derived from promoter sequences may also be biased to match those promoters. However, motifs derived from in vitro binding (e.g., [96]) avoid this potential bias or circularity.

This phylogenetic average motif score representation building approach does not use deep learning. However, could we improve the representation with deep learning? Recently, self-supervised training of natural language models on biological sequences and subsequent fine-tuning for specific tasks has led to new approaches for many classic problems in protein [97–99] and DNA[28] sequence analysis. Representation learning has the potential to learn motifs that are not found in the databases[27,100,101], as well as more complicated regulation and rules about motif spacing and orientation [94,95,102]; here we simply took the best match in each sequence. Because it uses no deep learning or trainable parameters and is based on freely available data, phylogenetic average motif score representations serve as a baseline for future representation learning efforts.

Taken together, the success of the phylogenetic average motif score representation to group together promoters and 3’UTRs with similar function indicates that combinations of transcription factor and RNA binding motifs may be sufficient to predict (at least some) functions for these non-coding regions. Finally, our results suggest that, at least when averaged over evolution, the “rules” and “grammar” of the cis-regulatory code may be simpler than is currently anticipated[1], but that global, high-dimensional views may be essential to its understanding.

## Methods

### Data collection of yeast and human non-coding sequences

The ideal choice of species for our approach would be those with maximum sequence divergence and minimal functional divergence; too close to the reference species and the promoters may be too similar, but too diverged and the TFs and gene expression patterns may not be conserved. For *S. cerevisiae*, we decided on up to 35 species per gene (Shen et al., 2016;[54]). In alphabetical order: *Ashbya gossypii, Candida albicans, Candida auris, Candida glabrata, Candida parapsilosis, Candida tropicalis, Clavispora lusitaniae, Debaryomyces fabryi, Debaryomyces hansenii, Eremothecium cymbalariae, Eremothecium gossypii, Kazachstania africana, Kazachstania naganishii, Kluyveromyces dobzhanskii, Kluyveromyces lactis, Kluyveromyces marxianus, Komagataella pastoris, Komagataella phaffii, Lachancea dasiensis, Lachancea lanzarotensis, Lodderomyces elongisporus, Naumovozyma castellii, Naumovozyma dairenensis, Rhodotorula graminis, Saccharomyces arboricola, Saccharomyces eubayanus, Saccharomyces boulardii, Saccharomycetaceae ashbya aceri, Tetrapisispora blattae, Tetrapisispora phaffii, Tortispora caseinolytica, Torulaspora delbrueckii, Vanderwaltozyma polyspora, Yarrowia lipolytica, and Zygosaccharomyces rouxii*. 6600 S. cerevisiae promoter sequences and up to 35 one-to-one orthologs per *S. cerevisiae* gene were retrieved from Ensembl[60]. They were filtered for only those sequences that contained the letters A, T, G, and C. Only those *S. cerevisiae* genes that were associated with a minimum of 10 orthologous species were used. The final representation for yeast promoter sequences contained 4337 *S. cerevisiae* genes with a total of 126,049 promoter sequences. 3’UTR sequences were obtained similarly, with 4294 *S. cerevisiae* genes in the representation and 82,187 3’UTR sequences in total. Human gene IDs for 18,256 genes were obtained from Ensembl and 2,612,699 homologous sequences were obtained from a Zoonomia alignment[103]. We then used a heuristic filter similar to the approach described elsewhere[101] to make the number of homologs more comparable to the yeast dataset. After this filtering, we were left with 814,963 sequences for the 18,256 promoters. We then applied the filters described above for the yeast sequences and obtained 800,178 sequences in 15,906 human homologous promoter sets.

### Collection of motif data

244 Yeast TF motifs were downloaded from YeTFaSCo[34] (expert curated motifs), 137 human TF motifs were downloaded from JASPAR[33] (non-redundant core vertebrate set), and 74 yeast RBP motifs were downloaded from AtTract[35]. All position frequency matrices were converted to PWMs using a log ratio of motif to a background frequency of 0.25 per nucleotide. For matrices that are given as nucleotide counts, a pseudo-count of 1 per position was added for smoothing. For matrices that were given as probabilities between 0 and 1, a heuristic procedure was used for smoothing. The effective number of observations, Ne, was defined as Ne = 1/min(p) where min(p) was the smallest non-zero number in the PFM probability matrix. The counts for nucleotide b and position i, were then estimated as c = Ne x p, where p is the probability of nucleotide b at position i. This estimated counts matrix was then softened with the usual 1 per position pseudo-count.

### Convolutional scanning and normalization to build the representation

To build the representation, we took advantage of GPUs and the speed of machine learning tools and used a convolutional layer in Keras/Tensorflow to compute match scores of motifs to promoter sequences. Motifs were embedded in a single keras convolutional layer with kernel size set to the largest motif (for yeast TF motifs, kernel size was 29; for human, kernel size was 35; for yeast RBP motifs, kernel size was 12). Motifs were centered in an array of 0s so that they were all a standard size for loading into a convolutional layer.

For promoter sequences, we embedded each motif and its reverse complement in a one-dimensional convolutional layer and scanned the motifs across the entire sequence. The greater of the two match scores was reported to be the match score for that position by performing a max pool over the forward and reverse scores. For 3’UTR sequences, since they are single-stranded as RNA, we only scanned forward motifs. Generally, a potential binding site is identified when the score of the PWM model is above an arbitrary threshold, and higher scores are interpreted as binding sites with greater strength than those with lower scores[2]. Thus, we returned the highest score of the PWM scanning for a particular sequence as the “best match” of the motif to that sequence by performing a max pool across the length of the sequence[2]. These “best matches” were averaged over all the available homologs for each promoter or 3’ UTR. We normalized these phylogenetic averaged “best match” scores by scrambling the motifs 100 times and repeating the convolutional scanning for each scrambled matrix, then computing a z-score between the real match scores and the distribution of the scrambled match scores.

To visualize the feature space, we performed hierarchical clustering on the data using Cluster 3.0[104] (median centered arrays, uncentered correlation on genes and arrays, average linkage) and visualized the clustering using Java Treeview[105]. Clusters were extracted from the feature space using Java Treeview[105]. Yeast clusters were annotated using the Saccharomyces Genome Database (SGD) GO term finder[61], and human clusters were annotated using g:Profiler[106]. Process, function, and component GO terms were considered. The false discovery rate was set to 0.05 and the terms were Benjamini-Hochberg corrected.

P-value threshold was set to 0.05.

### Collection and processing of gene expression and binding data

Gene expression data for yeast was obtained from a collection of stress experiments, where yeast were subjected to a large variety of growth conditions [63] (arrays were median centered) and a large set of genetic perturbations [64] (genes and arrays were median centered). Single cell type data[66] for human was obtained from The Human Protein Atlas and both genes and cell types were median centered. Median TPM RNA expression for tissues was obtained from the GTEx v8 portal. The median values for each gene were log transformed and then used as a measure of absolute expression level. The GTEx data were also log transformed and genes were median centered. A matrix of yeast promoters with direct evidence for binding of any transcription factor was retrieved from yeastract on July 26th, 2022.

### Nearest neighbour (NN) regression

To evaluate the predictive power of the feature space and compare between representations easily, we performed a nearest neighbour (NN) regression (also known as k-nearest neighbour with k=1) using a cosine similarity metric. Only those genes with less than 15% missing data points were included. For the remaining genes, the missing data was replaced with 0s. Nearest Neighbour regression predicts each point (in our case expression measurement) to be the value of its nearest neighbour (in our case in the sequence-based representation space). We measured the power of the regression for each set of genome-wide expression measurements using the Pearson correlation. We then summarized the Pearson correlations using the mean and standard error over all the experiments in the dataset. For comparison, a randomized Pearson correlation was calculated by shuffling the gene identifiers of the expression measurements and performing NN regression with the sequence-based representation space. Additionally, an upper bound Pearson correlation was determined by performing NN regression on the expression measurements against themselves.

### Sequence similarity of yeast and human promoter sequences

To calculate sequence similarity of promoter sequences in presented clusters, we used the Clustal Omega[107]. This generates a matrix of sequence similarity between all promoter sequences. To generate an expectation for what constitutes as low sequence similarity, we generated 100 random 500 bp sequences composed of an equal proportion of A, T, C, and G nucleotides and generated a matrix of sequence similarity with the random sequences. To get the average and standard deviation of the percent identity of these sequences, we averaged all the percent identities in the matrix except for the diagonal of 100% identities corresponding to the sequence similarity between a promoter and itself. The average and standard deviation of this percent identity matrix generated using random sequence was 28% ± 3% but the maximum percent identity was approximately 46%, while the minimum percent identity was approximately 19%. Taking this into account, and the fact that the nucleotides do not appear at a rate of 25% each in yeast or human sequences, we assumed that promoters in clusters presented had relatively low sequence similarity if the sequence similarity of most of the promoter pairs in the matrix was below 50%.

### Sequence similarity of nucleotide and protein sequences as baselines

To compare representations in the evolutionary analysis to known baselines, we used blastn [108] to find nearest neighbours of promoter sequences, and blastp [108] to find nearest neighbours of protein sequences (also retrieved from Ensembl). Starting with promoters, we first created a database of all *S. cerevisiae* promoter sequences, and then we performed “all-by-all” blast (maximum target sequences = 2, maximum e-value = 10) between the database and all the input sequences. We used the closest BLAST hit (other than itself), defined as the hit with the highest percent identity and lowest e-value, for a particular target sequence as its “neighbour”. We used these for nearest neighbour regression as described above. These steps were repeated for protein sequences.

### Building a representation for evolutionary analysis of TFs in yeast

To generate a representation for evolutionary analysis of TFs in yeast (Figure 6), we modified the encoder so that instead of averaging PWM max scores over all orthologs of a particular gene, we averaged PWM max scores over all genes in a particular cluster (for example, ribosome biosynthesis) of a particular species (for example, *C. albicans*).

## Acknowledgements

We thank Dr. Tim Hughes for the suggestion to display gene expression data. We thank Dr. Alex Lu for help with keras, GPUs and for comments on the manuscript. We thank Ami Sangster and Drs. Nirvana Nursimulu and Audrey Gasch for comments on the manuscript. The Genotype-Tissue Expression (GTEx) Project was supported by the Common Fund of the Office of the Director of the National Institutes of Health, and by NCI, NHGRI, NHLBI, NIDA, NIMH, and NINDS.

## Supplementary Figures

**Supplementary Figure S1.**
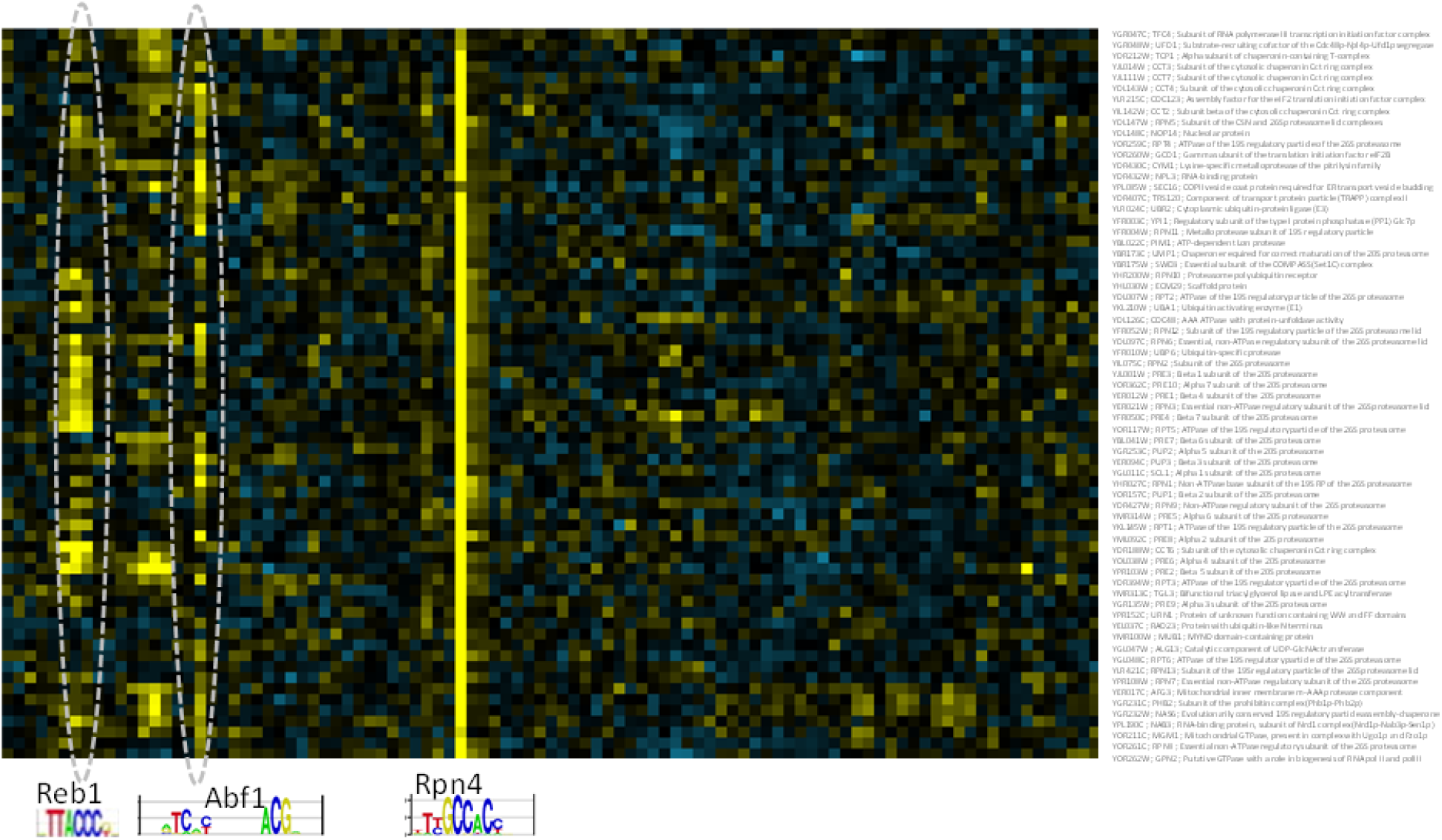
Abf1 or Reb1 are found with Rpn4 (Table 1, cluster G) in proteasomal promoters and the cytosolic chaperonin ring complex. Not all 244 yeast motifs are shown for clarity. Z-scores are displayed as in Figure 3

## Supplementary Data

Includes the raw phylogenetic average motif score representations as well as the files needed for the visualizations in figures 2 and 5. Also includes Excel spreadsheets with the information about each manually identified cluster and the enrichment analysis results shown in tables 1, 2 and 6.

## References

1. Zeitlinger J. Seven myths of how transcription factors read the cis-regulatory code. Curr Opin Syst Biol. 2020;23: 22–31. doi:10.1016/j.coisb.2020.08.002

2. Aerts S. Computational strategies for the genome-wide identification of cis-regulatory elements and transcriptional targets. Curr Top Dev Biol. 2012;98: 121–145. doi:10.1016/B978-0-12-386499-4.00005-7

3. Kim S, Wysocka J. Deciphering the multi-scale, quantitative cis-regulatory code. Mol Cell. 2023; S1097–2765(22)01215–1. doi:10.1016/j.molcel.2022.12.032

4. Barski A, Cuddapah S, Cui K, Roh T-Y, Schones DE, Wang Z, et al. High-resolution profiling of histone methylations in the human genome. Cell. 2007;129: 823–837. doi:10.1016/j.cell.2007.05.009

5. Rhee HS, Pugh BF. Comprehensive genome-wide protein-DNA interactions detected at single-nucleotide resolution. Cell. 2011;147: 1408–1419. doi:10.1016/j.cell.2011.11.013

6. Wang H, Johnston M, Mitra RD. Calling cards for DNA-binding proteins. Genome Res. 2007;17: 1202–1209. doi:10.1101/gr.6510207

7. Rossi MJ, Kuntala PK, Lai WKM, Yamada N, Badjatia N, Mittal C, et al. A high-resolution protein architecture of the budding yeast genome. Nature. 2021;592: 309–314. doi:10.1038/s41586-021-03314-8

8. ENCODE Project Consortium. An integrated encyclopedia of DNA elements in the human genome. Nature. 2012;489: 57–74. doi:10.1038/nature11247

9. Barrett LW, Fletcher S, Wilton SD. Regulation of eukaryotic gene expression by the untranslated gene regions and other non-coding elements. Cell Mol Life Sci CMLS. 2012;69: 3613–3634. doi:10.1007/s00018-012-0990-9

10. Barash Y, Calarco JA, Gao W, Pan Q, Wang X, Shai O, et al. Deciphering the splicing code. Nature. 2010;465: 53–59. doi:10.1038/nature09000

11. Lu Z, Guan X, Schmidt CA, Matera AG. RIP-seq analysis of eukaryotic Sm proteins identifies three major categories of Sm-containing ribonucleoproteins. Genome Biol. 2014;15: R7. doi:10.1186/gb-2014-15-1-r7

12. Van Nostrand EL, Pratt GA, Shishkin AA, Gelboin-Burkhart C, Fang MY, Sundararaman B, et al. Robust transcriptome-wide discovery of RNA-binding protein binding sites with enhanced CLIP (eCLIP). Nat Methods. 2016;13: 508–514. doi:10.1038/nmeth.3810

13. Kang Y, Patel NR, Shively C, Recio PS, Chen X, Wranik BJ, et al. Dual threshold optimization and network inference reveal convergent evidence from TF binding locations and TF perturbation responses. Genome Res. 2020;30: 459–471. doi:10.1101/gr.259655.119

14. Panchy NL, Lloyd JP, Shiu S-H. Improved recovery of cell-cycle gene expression in Saccharomyces cerevisiae from regulatory interactions in multiple omics data. BMC Genomics. 2020;21: 159. doi:10.1186/s12864-020-6554-8

15. Kelley DR, Snoek J, Rinn JL. Basset: learning the regulatory code of the accessible genome with deep convolutional neural networks. Genome Res. 2016;26: 990–999. doi:10.1101/gr.200535.115

16. Kelley DR, Reshef YA, Bileschi M, Belanger D, McLean CY, Snoek J. Sequential regulatory activity prediction across chromosomes with convolutional neural networks. Genome Res. 2018;28: 739–750. doi:10.1101/gr.227819.117

17. Avsec Ž, Agarwal V, Visentin D, Ledsam JR, Grabska-Barwinska A, Taylor KR, et al. Effective gene expression prediction from sequence by integrating long-range interactions. Nat Methods. 2021;18: 1196–1203. doi:10.1038/s41592-021-01252-x

18. de Boer CG, Vaishnav ED, Sadeh R, Abeyta EL, Friedman N, Regev A. Deciphering eukaryotic gene-regulatory logic with 100 million random promoters. Nat Biotechnol. 2020;38: 56–65. doi:10.1038/s41587-019-0315-8

19. Taskiran II, Spanier KI, Christiaens V, Mauduit D, Aerts S. Cell type directed design of synthetic enhancers. bioRxiv. 2022; 2022–07.

20. Vaishnav ED, de Boer CG, Molinet J, Yassour M, Fan L, Adiconis X, et al. The evolution, evolvability and engineering of gene regulatory DNA. Nature. 2022;603: 455–463. doi:10.1038/s41586-022-04506-6

21. Schreiber J, Singh R, Bilmes J, Noble WS. A pitfall for machine learning methods aiming to predict across cell types. Genome Biol. 2020;21: 282. doi:10.1186/s13059-020-02177-y

22. Karollus A, Mauermeier T, Gagneur J. Current sequence-based models capture gene expression determinants in promoters but mostly ignore distal enhancers. Genome Biol. 2023;24: 56. doi:10.1186/s13059-023-02899-9

23. Matsoukas C, Haslum JF, Sorkhei M, Söderberg M, Smith K. What Makes Transfer Learning Work For Medical Images: Feature Reuse & Other Factors. arXiv; 2022. doi:10.48550/arXiv.2203.01825

24. Neyshabur B, Sedghi H, Zhang C. What is being transferred in transfer learning? arXiv; 2021. doi:10.48550/arXiv.2008.11687

25. Raghu M, Zhang C, Kleinberg J, Bengio S. Transfusion: Understanding Transfer Learning for Medical Imaging. arXiv; 2019. doi:10.48550/arXiv.1902.07208

26. Lu AX, Lu AX, Schormann W, Ghassemi M, Andrews DW, Moses AM. The Cells Out of Sample (COOS) dataset and benchmarks for measuring out-of-sample generalization of image classifiers. arXiv; 2020. doi:10.48550/arXiv.1906.07282

27. Koo PK, Eddy SR. Representation learning of genomic sequence motifs with convolutional neural networks. PLOS Comput Biol. 2019;15: e1007560. doi:10.1371/journal.pcbi.1007560

28. Ji Y, Zhou Z, Liu H, Davuluri RV. DNABERT: pre-trained Bidirectional Encoder Representations from Transformers model for DNA-language in genome. Bioinformatics. 2021;37: 2112–2120. doi:10.1093/bioinformatics/btab083

29. Singh G, Mullany S, Moorthy SD, Zhang R, Mehdi T, Tian R, et al. A flexible repertoire of transcription factor binding sites and a diversity threshold determines enhancer activity in embryonic stem cells. Genome Res. 2021;31: 564–575. doi:10.1101/gr.272468.120

30. Novakovsky G, Fornes O, Saraswat M, Mostafavi S, Wasserman WW. ExplaiNN: interpretable and transparent neural networks for genomics. bioRxiv; 2022. p. 2022.05.20.492818. doi:10.1101/2022.05.20.492818

31. Yuan Y, Guo L, Shen L, Liu JS. Predicting Gene Expression from Sequence: A Reexamination. PLOS Comput Biol. 2007;3: e243. doi:10.1371/journal.pcbi.0030243

32. Zerbino DR, Achuthan P, Akanni W, Amode MR, Barrell D, Bhai J, et al. Ensembl 2018. Nucleic Acids Res. 2018;46: D754–D761. doi:10.1093/nar/gkx1098

33. Castro-Mondragon JA, Riudavets-Puig R, Rauluseviciute I, Lemma RB, Turchi L, Blanc-Mathieu R, et al. JASPAR 2022: the 9th release of the open-access database of transcription factor binding profiles. Nucleic Acids Res. 2022;50: D165–D173. doi:10.1093/nar/gkab1113

34. de Boer CG, Hughes TR. YeTFaSCo: a database of evaluated yeast transcription factor sequence specificities. Nucleic Acids Res. 2012;40: D169–179. doi:10.1093/nar/gkr993

35. Giudice G, Sánchez-Cabo F, Torroja C, Lara-Pezzi E. ATtRACT-a database of RNA-binding proteins and associated motifs. Database J Biol Databases Curation. 2016;2016: baw035. doi:10.1093/database/baw035

36. Ray D, Kazan H, Cook KB, Weirauch MT, Najafabadi HS, Li X, et al. A compendium of RNA-binding motifs for decoding gene regulation. Nature. 2013;499: 172–177. doi:10.1038/nature12311

37. Weirauch MT, Yang A, Albu M, Cote AG, Montenegro-Montero A, Drewe P, et al. Determination and Inference of Eukaryotic Transcription Factor Sequence Specificity. Cell. 2014;158: 1431–1443. doi:10.1016/j.cell.2014.08.009

38. Wasserman WW, Sandelin A. Applied bioinformatics for the identification of regulatory elements. Nat Rev Genet. 2004;5: 276–287. doi:10.1038/nrg1315

39. Glenwinkel L, Wu D, Minevich G, Hobert O. TargetOrtho: a phylogenetic footprinting tool to identify transcription factor targets. Genetics. 2014;197: 61–76. doi:10.1534/genetics.113.160721

40. Moses AM, Chiang DY, Pollard DA, Iyer VN, Eisen MB. MONKEY: identifying conserved transcription-factor binding sites in multiple alignments using a binding site-specific evolutionary model. Genome Biol. 2004;5: R98. doi:10.1186/gb-2004-5-12-r98

41. Arnold P, Erb I, Pachkov M, Molina N, van Nimwegen E. MotEvo: integrated Bayesian probabilistic methods for inferring regulatory sites and motifs on multiple alignments of DNA sequences. Bioinforma Oxf Engl. 2012;28: 487–494. doi:10.1093/bioinformatics/btr695

42. Pollard DA, Moses AM, Iyer VN, Eisen MB. Detecting the limits of regulatory element conservation and divergence estimation using pairwise and multiple alignments. BMC Bioinformatics. 2006;7: 376. doi:10.1186/1471-2105-7-376

43. Moses AM, Pollard DA, Nix DA, Iyer VN, Li X-Y, Biggin MD, et al. Large-scale turnover of functional transcription factor binding sites in Drosophila. PLoS Comput Biol. 2006;2: e130. doi:10.1371/journal.pcbi.0020130

44. Bradley RK, Li X-Y, Trapnell C, Davidson S, Pachter L, Chu HC, et al. Binding site turnover produces pervasive quantitative changes in transcription factor binding between closely related Drosophila species. PLoS Biol. 2010;8: e1000343. doi:10.1371/journal.pbio.1000343

45. He BZ, Holloway AK, Maerkl SJ, Kreitman M. Does positive selection drive transcription factor binding site turnover? A test with Drosophila cis-regulatory modules. PLoS Genet. 2011;7: e1002053. doi:10.1371/journal.pgen.1002053

46. Dermitzakis ET, Clark AG. Evolution of transcription factor binding sites in Mammalian gene regulatory regions: conservation and turnover. Mol Biol Evol. 2002;19: 1114–1121. doi:10.1093/oxfordjournals.molbev.a004169

47. Krieger G, Lupo O, Wittkopp P, Barkai N. Evolution of transcription factor binding through sequence variations and turnover of binding sites. Genome Res. 2022;32: 1099–1111. doi:10.1101/gr.276715.122

48. Bininda-Emonds OR, Brady SG, Kim J, Sanderson MJ. Scaling of accuracy in extremely large phylogenetic trees. Pac Symp Biocomput Pac Symp Biocomput. 2001; 547–558. doi:10.1142/9789814447362_0053

49. Price MN, Dehal PS, Arkin AP. FastTree: Computing Large Minimum Evolution Trees with Profiles instead of a Distance Matrix. Mol Biol Evol. 2009;26: 1641–1650. doi:10.1093/molbev/msp077

50. Wittkopp PJ, Kalay G. Cis-regulatory elements: molecular mechanisms and evolutionary processes underlying divergence. Nat Rev Genet. 2011;13: 59–69. doi:10.1038/nrg3095

51. Johnson AD. The rewiring of transcription circuits in evolution. Curr Opin Genet Dev. 2017;47: 121–127. doi:10.1016/j.gde.2017.09.004

52. Gasch AP, Moses AM, Chiang DY, Fraser HB, Berardini M, Eisen MB. Conservation and evolution of cis-regulatory systems in ascomycete fungi. PLoS Biol. 2004;2: e398. doi:10.1371/journal.pbio.0020398

53. Hogan GJ, Brown PO, Herschlag D. Evolutionary Conservation and Diversification of Puf RNA Binding Proteins and Their mRNA Targets. PLoS Biol. 2015;13: e1002307. doi:10.1371/journal.pbio.1002307

54. Wilinski D, Buter N, Klocko AD, Lapointe CP, Selker EU, Gasch AP, et al. Recurrent rewiring and emergence of RNA regulatory networks. Proc Natl Acad Sci U S A. 2017;114: E2816–E2825. doi:10.1073/pnas.1617777114

55. Balcı AT, Ebeid MM, Benos PV, Kostka D, Chikina M. An intrinsically interpretable neural network architecture for sequence to function learning. bioRxiv; 2023. p. 2023.01.25.525572. doi:10.1101/2023.01.25.525572

56. Storcheus D, Rostamizadeh A, Kumar S. A Survey of Modern Questions and Challenges in Feature Extraction. Proceedings of the 1st International Workshop on Feature Extraction: Modern Questions and Challenges at NIPS 2015. PMLR; 2015. pp. 1–18. Available: https://proceedings.mlr.press/v44/storcheus2015survey.html

57. Xu T, Zheng X, Li B, Jin P, Qin Z, Wu H. A comprehensive review of computational prediction of genome-wide features. Brief Bioinform. 2020;21: 120–134. doi:10.1093/bib/bby110

58. Bonidia RP, Sampaio LDH, Domingues DS, Paschoal AR, Lopes FM, de Carvalho ACPLF, et al. Feature extraction approaches for biological sequences: a comparative study of mathematical features. Brief Bioinform. 2021;22: bbab011. doi:10.1093/bib/bbab011

59. Boy-Marcotte E, Lagniel G, Perrot M, Bussereau F, Boudsocq A, Jacquet M, et al. The heat shock response in yeast: differential regulations and contributions of the Msn2p/Msn4p and Hsf1p regulons. Mol Microbiol. 1999;33: 274–283. doi:10.1046/j.1365-2958.1999.01467.x

60. Cunningham F, Allen JE, Allen J, Alvarez-Jarreta J, Amode MR, Armean IM, et al. Ensembl 2022. Nucleic Acids Res. 2022;50: D988–D995. doi:10.1093/nar/gkab1049

61. Cherry JM, Hong EL, Amundsen C, Balakrishnan R, Binkley G, Chan ET, et al. Saccharomyces Genome Database: the genomics resource of budding yeast. Nucleic Acids Res. 2012;40: D700–705. doi:10.1093/nar/gkr1029

62. Eisen MB, Spellman PT, Brown PO, Botstein D. Cluster analysis and display of genome-wide expression patterns. Proc Natl Acad Sci U S A. 1998;95: 14863–14868. doi:10.1073/pnas.95.25.14863

63. Talavera D, Kershaw CJ, Costello JL, Castelli LM, Rowe W, Sims PFG, et al. Archetypal transcriptional blocks underpin yeast gene regulation in response to changes in growth conditions. Sci Rep. 2018;8: 7949. doi:10.1038/s41598-018-26170-5

64. Kemmeren P, Sameith K, van de Pasch LAL, Benschop JJ, Lenstra TL, Margaritis T, et al. Large-Scale Genetic Perturbations Reveal Regulatory Networks and an Abundance of Gene-Specific Repressors. Cell. 2014;157: 740–752. doi:10.1016/j.cell.2014.02.054

65. Gasch AP, Spellman PT, Kao CM, Carmel-Harel O, Eisen MB, Storz G, et al. Genomic expression programs in the response of yeast cells to environmental changes. Mol Biol Cell. 2000;11: 4241–4257. doi:10.1091/mbc.11.12.4241

66. Karlsson M, Zhang C, Méar L, Zhong W, Digre A, Katona B, et al. A single-cell type transcriptomics map of human tissues. Sci Adv. 2021;7: eabh2169. doi:10.1126/sciadv.abh2169

67. Thiel G, Ekici M, Rössler OG. RE-1 silencing transcription factor (REST): a regulator of neuronal development and neuronal/endocrine function. Cell Tissue Res. 2015;359: 99–109. doi:10.1007/s00441-014-1963-0

68. Zhou Q, Yu M, Tirado-Magallanes R, Li B, Kong L, Guo M, et al. ZNF143 mediates CTCF-bound promoter–enhancer loops required for murine hematopoietic stem and progenitor cell function. Nat Commun. 2021;12: 43. doi:10.1038/s41467-020-20282-1

69. Beagan JA, Duong MT, Titus KR, Zhou L, Cao Z, Ma J, et al. YY1 and CTCF orchestrate a 3D chromatin looping switch during early neural lineage commitment. Genome Res. 2017;27: 1139–1152. doi:10.1101/gr.215160.116

70. Capps D, Hunter A, Chiang M, Pracheil T, Liu Z. Ubiquitin-Conjugating Enzymes Ubc1 and Ubc4 Mediate the Turnover of Hap4, a Master Regulator of Mitochondrial Biogenesis in Saccharomyces cerevisiae. Microorganisms. 2022;10: 2370. doi:10.3390/microorganisms10122370

71. Ramos F, Dubois E, Piérard A. Control of enzyme synthesis in the lysine biosynthetic pathway of Saccharomyces cerevisiae. Evidence for a regulatory role of gene LYS14. Eur J Biochem. 1988;171: 171–176. doi:10.1111/j.1432-1033.1988.tb13773.x

72. Hinnebusch AG, Natarajan K. Gcn4p, a master regulator of gene expression, is controlled at multiple levels by diverse signals of starvation and stress. Eukaryot Cell. 2002;1: 22–32. doi:10.1128/EC.01.1.22-32.2002

73. Kumar R, Reynolds DM, Shevchenko A, Shevchenko A, Goldstone SD, Dalton S. Forkhead transcription factors, Fkh1p and Fkh2p, collaborate with Mcm1p to control transcription required for M-phase. Curr Biol CB. 2000;10: 896–906. doi:10.1016/s0960-9822(00)00618-7

74. Pic A, Lim FL, Ross SJ, Veal EA, Johnson AL, Sultan MR, et al. The forkhead protein Fkh2 is a component of the yeast cell cycle transcription factor SFF. EMBO J. 2000;19: 3750–3761. doi:10.1093/emboj/19.14.3750

75. Zhu G, Davis TN. The fork head transcription factor Hcm1p participates in the regulation of SPC110, which encodes the calmodulin-binding protein in the yeast spindle pole body. Biochim Biophys Acta. 1998;1448: 236–244. doi:10.1016/s0167-4889(98)00135-9

76. Horak CE, Luscombe NM, Qian J, Bertone P, Piccirrillo S, Gerstein M, et al. Complex transcriptional circuitry at the G1/S transition in Saccharomyces cerevisiae. Genes Dev. 2002;16: 3017–3033. doi:10.1101/gad.1039602

77. Mannhaupt G, Schnall R, Karpov V, Vetter I, Feldmann H. Rpn4p acts as a transcription factor by binding to PACE, a nonamer box found upstream of 26S proteasomal and other genes in yeast. FEBS Lett. 1999;450: 27–34. doi:10.1016/s0014-5793(99)00467-6

78. Remacle JE, Holmberg S. A REB1-binding site is required for GCN4-independent ILV1 basal level transcription and can be functionally replaced by an ABF1-binding site. Mol Cell Biol. 1992;12: 5516–5526. doi:10.1128/mcb.12.12.5516-5526.1992

79. Belaguli NS, Schildmeyer LA, Schwartz RJ. Organization and myogenic restricted expression of the murine serum response factor gene. A role for autoregulation. J Biol Chem. 1997;272: 18222–18231. doi:10.1074/jbc.272.29.18222

80. UniProt Consortium. UniProt: the universal protein knowledgebase in 2021. Nucleic Acids Res. 2021;49: D480–D489. doi:10.1093/nar/gkaa1100

81. Yeang C-H, Mak HC, McCuine S, Workman C, Jaakkola T, Ideker T. Validation and refinement of gene-regulatory pathways on a network of physical interactions. Genome Biol. 2005;6: R62. doi:10.1186/gb-2005-6-7-r62

82. Wang L, Hou L, Qian M, Li F, Deng M. Integrating multiple types of data to predict novel cell cycle-related genes. BMC Syst Biol. 2011;5 Suppl 1: S9. doi:10.1186/1752-0509-5-S1-S9

83. Williams RM, Primig M, Washburn BK, Winzeler EA, Bellis M, Sarrauste de Menthiere C, et al. The Ume6 regulon coordinates metabolic and meiotic gene expression in yeast. Proc Natl Acad Sci U S A. 2002;99: 13431–13436. doi:10.1073/pnas.202495299

84. Dreyfuss G, Kim VN, Kataoka N. Messenger-RNA-binding proteins and the messages they carry. Nat Rev Mol Cell Biol. 2002;3: 195–205. doi:10.1038/nrm760

85. Gerber AP, Herschlag D, Brown PO. Extensive association of functionally and cytotopically related mRNAs with Puf family RNA-binding proteins in yeast. PLoS Biol. 2004;2: E79. doi:10.1371/journal.pbio.0020079

86. Tanay A, Regev A, Shamir R. Conservation and evolvability in regulatory networks: the evolution of ribosomal regulation in yeast. Proc Natl Acad Sci U S A. 2005;102: 7203–7208. doi:10.1073/pnas.0502521102

87. Mallick J, Whiteway M. The evolutionary rewiring of the ribosomal protein transcription pathway modifies the interaction of transcription factor heteromer Ifh1-Fhl1 (interacts with forkhead 1-forkhead-like 1) with the DNA-binding specificity element. J Biol Chem. 2013;288: 17508–17519. doi:10.1074/jbc.M112.436683

88. Hogues H, Lavoie H, Sellam A, Mangos M, Roemer T, Purisima E, et al. Transcription factor substitution during the evolution of fungal ribosome regulation. Mol Cell. 2008;29: 552–562. doi:10.1016/j.molcel.2008.02.006

89. Lavoie H, Hogues H, Mallick J, Sellam A, Nantel A, Whiteway M. Evolutionary tinkering with conserved components of a transcriptional regulatory network. PLoS Biol. 2010;8: e1000329. doi:10.1371/journal.pbio.1000329

90. Sarda S, Hannenhalli S. High-Throughput Identification of Cis-Regulatory Rewiring Events in Yeast. Mol Biol Evol. 2015;32: 3047–3063. doi:10.1093/molbev/msv203

91. Huber A, French SL, Tekotte H, Yerlikaya S, Stahl M, Perepelkina MP, et al. Sch9 regulates ribosome biogenesis via Stb3, Dot6 and Tod6 and the histone deacetylase complex RPD3L. EMBO J. 2011;30: 3052–3064. doi:10.1038/emboj.2011.221

92. Liko D, Slattery MG, Heideman W. Stb3 binds to ribosomal RNA processing element motifs that control transcriptional responses to growth in Saccharomyces cerevisiae. J Biol Chem. 2007;282: 26623–26628. doi:10.1074/jbc.M704762200

93. Teixeira MC, Monteiro PT, Palma M, Costa C, Godinho CP, Pais P, et al. YEASTRACT: an upgraded database for the analysis of transcription regulatory networks in Saccharomyces cerevisiae. Nucleic Acids Res. 2018;46: D348–D353. doi:10.1093/nar/gkx842

94. Avsec Ž, Weilert M, Shrikumar A, Krueger S, Alexandari A, Dalal K, et al. Base-resolution models of transcription-factor binding reveal soft motif syntax. Nat Genet. 2021;53: 354–366. doi:10.1038/s41588-021-00782-6

95. de Almeida BP, Reiter F, Pagani M, Stark A. DeepSTARR predicts enhancer activity from DNA sequence and enables the de novo design of synthetic enhancers. Nat Genet. 2022;54: 613–624. doi:10.1038/s41588-022-01048-5

96. Pelossof R, Singh I, Yang JL, Weirauch MT, Hughes TR, Leslie CS. Affinity regression predicts the recognition code of nucleic acid-binding proteins. Nat Biotechnol. 2015;33: 1242–1249. doi:10.1038/nbt.3343

97. Rives A, Meier J, Sercu T, Goyal S, Lin Z, Liu J, et al. Biological structure and function emerge from scaling unsupervised learning to 250 million protein sequences. Proc Natl Acad Sci. 2021;118: e2016239118. doi:10.1073/pnas.2016239118

98. Stärk H, Dallago C, Heinzinger M, Rost B. Light attention predicts protein location from the language of life. Bioinforma Adv. 2021;1: vbab035. doi:10.1093/bioadv/vbab035

99. Chandra A, Tünnermann L, Löfstedt T, Gratz R. Transformer-based deep learning for predicting protein properties in the life sciences. Dötsch V, editor. eLife. 2023;12: e82819. doi:10.7554/eLife.82819

100. Koo PK, Ploenzke M. Deep learning for inferring transcription factor binding sites. Curr Opin Syst Biol. 2020;19: 16–23. doi:10.1016/j.coisb.2020.04.001

101. Lu AX, Lu AX, Pritišanac I, Zarin T, Forman-Kay JD, Moses AM. Discovering molecular features of intrinsically disordered regions by using evolution for contrastive learning. PLoS Comput Biol. 2022;18: e1010238. doi:10.1371/journal.pcbi.1010238

102. Chen L, Capra JA. Learning and interpreting the gene regulatory grammar in a deep learning framework. PLoS Comput Biol. 2020;16: e1008334. doi:10.1371/journal.pcbi.1008334

103. Zoonomia Consortium. A comparative genomics multitool for scientific discovery and conservation. Nature. 2020;587: 240–245. doi:10.1038/s41586-020-2876-6

104. de Hoon MJL, Imoto S, Nolan J, Miyano S. Open source clustering software. Bioinforma Oxf Engl. 2004;20: 1453–1454. doi:10.1093/bioinformatics/bth078

105. Saldanha AJ. Java Treeview—extensible visualization of microarray data. Bioinformatics. 2004;20: 3246–3248. doi:10.1093/bioinformatics/bth349

106. Raudvere U, Kolberg L, Kuzmin I, Arak T, Adler P, Peterson H, et al. g:Profiler: a web server for functional enrichment analysis and conversions of gene lists (2019 update). Nucleic Acids Res. 2019;47: W191–W198. doi:10.1093/nar/gkz369

107. Sievers F, Higgins DG. Clustal omega. Curr Protoc Bioinforma. 2014;48:3.13.1-3.13.16. doi:10.1002/0471250953.bi0313s48

108. Altschul SF, Madden TL, Schäffer AA, Zhang J, Zhang Z, Miller W, et al. Gapped BLAST and PSI-BLAST: a new generation of protein database search programs. Nucleic Acids Res. 1997;25: 3389–3402. doi:10.1093/nar/25.17.3389

